# *In vitro* EAS-mediated activity of *Alternaria* toxins

**DOI:** 10.64898/2026.07.09.737498

**Authors:** Eliana Spilioti, Anastasia Spyropoulou, Laurent Gaté, Mylène Lorcin, Kyriaki Machera, Alexandra Nestora, Anastasia Repouskou, Ioannis Theologidis, Doris Marco, Anne-Cathrin Behr

## Abstract

*Alternaria* mycotoxins represent an emerging concern due to their frequent occurrence in food and feed. However, available toxicological data remain limited. Under the current EU regulatory framework, and in line with the EFSA/ECHA/JRC guidance for the identification of endocrine disruptors (EDs), assessment of endocrine activity relies on standardized assays performed according to OECD Test Guidelines (TGs) for the estrogen-, androgen- and steroidogenesis- (EAS) modalities. Within the framework of the European Partnership for the Assessment of Risks from Chemicals (PARC), standardized *in vitro* methods of regulatory relevance were performed for six chemically characterized *Alternaria* toxins, aiming to address current regulatory gaps on EAS-mediated activity. Alternariol (AOH), alternariol monomethyl ether (AME), tenuazonic acid (TeA), altertoxin-I (ATX-I), tentoxin (TEN) and altenuene (ALT) were assessed over a broad concentration range, from 0.001 up to 60 μM, depending on cytotoxicity and solubility profile of each compound. Our findings indicate estrogenic activity for AOH (PC_50_: 3.9 - 4.6 µM) and AME (PC_50_: 5.2 - 8.5 µM) in the estrogen receptor transactivation assay (OECD TG 455), as well as an anti-estrogenic activity for ATX-I (IC_30_: 0.27 - 0.37 µM). Minimal positive responses were observed at high concentrations for AOH (from the dose of 3 µM) and for AME (from the dose of 10 µM) in the agonistic part of the androgen receptor transactivation assay (OECD TG 458), which may also reflect glucocorticoid receptor activation. No effects on estradiol or testosterone production were observed for any of the tested *Alternaria* compounds in the steroidogenesis assay (OECD TG 456).

## Introduction

Mycotoxins are secondary metabolites produced by various fungal species, with environmental conditions playing a key role in their production (Perrone et al. 2020). *Alternaria* mycotoxins are produced by the filamentous fungi of the genus *Alternaria.* They represent an emerging concern due to their frequent occurrence in food and feed (Zhang et al. 2025). The EFSA Panel on Contaminants in the Food Chain (CONTAM Panel) published the first risk assessment for *Alternaria* toxins in 2011, however due to insufficient toxicity data, health guidance values could not be established (EFSA 2011). Based on a more recent EFSA scientific report (2016), the estimated chronic dietary exposure to the *Alternaria* toxins alternariol (AOH), alternariol monomethyl ether (AME) and tenuazonic acid (TeA) was found to exceed the relevant threshold of toxicological concern (EFSA 2016). In 2022, the EU issued a Commission Recommendation on monitoring the occurrence of *Alternaria* toxins (EC 2022). An urgent need for additional compound-specific toxicity data is clearly indicated.

Although several *Alternaria* toxins including AOH, AME, TeA, altertoxin-I (ATX-I), tentoxin (TEN) and altenuene (ALT) have been chemically characterized, *in vivo* toxicological data remain limited. Most available evidence comes from *in vitro* studies, with cytotoxicity and genotoxicity being the most extensively investigated endpoints (Louro et al. 2024). Increasing attention has also been given to the potential of *Alternaria* toxins to disrupt the endocrine system. In the 2011 EFSA Scientific Opinion, *in vitro* mechanistic data were evaluated in the context of reproductive and developmental toxicity. Although it was concluded that AOH and AME may interfere with hormonal signaling pathways, no firm conclusion could be drawn due to limited and inconsistent data, and the underlying mechanisms were proposed to be cell-type specific and thus dependent on the type of the experimental cell model (EFSA 2011).

The endocrine-related activity of *Alternaria* toxins has been further investigated in a number of *in vitro* studies using different cell systems (Louro et al. 2024). Weak estrogenic activity has been reported for AOH in two studies using MMV-Luc cells (Demaegdt et al. 2016; Frizzell et al. 2013). AOH and AME have been also reported to consistently increase alkaline phosphatase (ALP) mRNA and/or enzymatic activity in Ishikawa cells, but the potency was as low as 0.01% of that of estradiol (E2) (Aichinger et al. 2020; Aichinger et al. 2019; Dellafiora et al. 2018; Lehmann et al. 2006). In the same cell line AOH has been shown to induce nuclear translocation of estrogen receptor (ER) α at the highest tested concentration of 10 µM [12]. *In silico* data indicate that AOH and AME can bind to both ERα and ERβ ligand pockets but with low relative binding affinity compared to estradiol (Dellafiora et al. 2018). Possible interactions with the androgen receptor (AR) have been also examined for AOH in two different studies using the androgen dependent TARM-Luc cell line and no agonistic effects were observed (Demaegdt et al. 2016; Frizzell et al. 2013). In a yeast androgen bioassay, AOH was reported to exhibit full agonistic activity, despite its relatively low potency (EC_50_: 270 μM) (Stypuła-Trębas et al. 2017). Weak antagonistic effects have been reported in TARM-Luc cells, however, findings should be interpreted cautiously as cytotoxicity was also observed in most of the corresponding doses when cell viability was assessed (Demaegdt et al. 2016; Frizzell et al. 2013).

Data on the effects of *Alternaria* toxins on steroidogenesis are scarce, with some indications for potential interference. In a study using the H295R adrenal cell line, AOH was shown to increase progesterone and estradiol synthesis and to induce mRNA expression of steroidogenesis-related genes (Frizzell et al. 2013). The authors further examined the proteomic response of H295R cells to AOH, and identified regulated proteins involved in the early-stage steroid biosynthesis and in C21-steroid hormone metabolism (Kalayou et al. 2014). On the other hand, AOH and AME administration to pig granulosa cells, decreased the level of progesterone secretion which was associated with reduced abundance of P450 cholesterol side-chain cleavage enzyme (P450scc), but not with changes in the mRNA expression of steroidogenesis-related genes. TeA showed no effect on progesterone secretion in the same cell system (Tiemann et al. 2009).

Overall, the available *in vitro* and *in silico* data demonstrate that *Alternaria* toxins may interfere with the endocrine system. Under the current EU regulatory framework, and in line with the EFSA/ECHA/JRC guidance, identification of Endocrine Disruptors (EDs) requires evidence for endocrine adversity at the organism level, endocrine activity and a biologically plausible link (EFSA 2018). The assessment of endocrine activity for the Estrogen, Androgen and Steroidogenesis (EAS)-modalities relies on standardized assays performed according to OECD Test Guidelines (TGs). These include the OECD TGs 455 and 458 for ER- and AR-mediated activity (OECD 2023), (OECD 2025b) and the OECD TG 456 for effects on E2 or testosterone (T) production (OECD 2025b). Within the framework of the European Partnership for the Assessment of Risks from Chemicals (PARC), standardized *in vitro* methods of regulatory relevance have been conducted for *Alternaria* toxins aiming to fill the gaps on EAS-mediated activity. The E-Screen cell proliferation assay in MCF-7 cells was also conducted, and results are presented as supplementary to the OECD TG 455.

## 2. Material and Methods

### 2.1 Alternaria toxins

Alternariol (AOH, CAS 641-38-3) and alternariol-9-methyl ether (AME, CAS 23452-05-3) were kindly synthesized by Prof. Süßmuth (TU Berlin, Germany). AOH was synthesized according to Won et al. (Won et al. 2015). The convergent synthesis featured a formylated aryl bromide and a desymmetrized aryl borane. Both were connected during a Pd-mediated Suzuki coupling. The final ring closure of the oxidized biaryl was performed with an ester formation during demethylation of all remaining alcohols. The product was isolated using reverse phase column chromatography and the identity of the material was confirmed by analytical HPLC-ESI-MS(MS) and ^1^HNMR spectroscopy. The convergent synthesis of AME was carried out according to Mikula et al. (Mikula et al. 2013). The building blocks were prepared from orcinol and 3,5-dimethoxyaniline respectively and coupled using diethyl chlorophosphate (DECP). The benzocoumarin core structure was established by intramolecular cyclization of the iodoresorcylic acid phenyl ester. Selective demethylation of alternariol 2,4-dimethyl ether (ADME) with AlCl3/NaI yielded AME which was isolated using reverse phase column chromatography. The identity of the material was confirmed by analytical HPLC-ESI-MS(MS) and ^1^H NMR spectroscopy.

Tenuazonic acid (TeA, CAS 610-88-8, purity > 95%) was purchased from Santa Cruz Biotechnology (Heidelberg, Germany), and altertoxin-I (ATX-I, CAS 56258-32-3, purity: 95.9%), (-)-altenuene (ALT, CAS 889101-41-1, purity: 99%) and tentoxin (TEN, CAS 28540-82-1, purity: 99.5%) were purchased from Biomol (Hamburg, Germany). All assays were conducted using the same batch of each mycotoxin to maximize data comparability and minimize uncertainties associated with batch effects. Upon arrival, dry compounds were stored at -80°C. The mycotoxins were dissolved in dimethyl sulfoxide (DMSO) at a maximum concentration of 10-30 mM. Stock aliquots were prepared in amber glass vials and more than three freeze-thaw cycles were avoided. Stock solutions were stored at -20°C.

### 2.2 Estrogen receptor stably transfected transactivation assay (ER STTA)

#### 2.2.1 Chemicals and reagents

E2 (CAS 50-28-2, purity ≥ 98%), 4-hydroxytamoxifen (OHT, CAS 68047-06-3, purity ≥ 98%) and cell culture grade DMSO (Cat. No D4540) were purchased from Sigma Aldrich (Saint Quentin Fallavier, France). Phenol-red-free Eagle’s minimum essential medium (MEM, Cat. No 10338343) and kanamycin (Cat. No 15160054) were obtained from Gibco (Grand Island, NY, USA). Dextran charcoal-stripped fetal bovine serum (Cat. No S181F-500) was purchased from Biowest (Nuaillé, France), WST-1 (Cat. No 11,644,807,001) from Roche (Manheim, Germany), Cell Culture Lysis reagent (Cat. No E1531) and One-Glo Luciferase (Cat. No E6120) were obtained from Promega (Charbonnières les Bains, France).

#### 2.2.2 Cell culture

The hERα-HeLa9903 cell line was purchased from the Japanese Collection of Research Bioresources Cell Bank (JCRB1318-HeLa9903, JCRB, Osaka Japan). The cell line can be used to identify human estrogen receptor α (hERα) agonists and antagonists, as detailed in the OECD test guideline 455 for the testing of chemicals (OECD 2018) (OECD, 2021). The hERα-HeLa9903 cell line is derived from the cervical tumor cells HeLa, stably transfected with a plasmid encoding human estrogen receptor α and a second plasmid with a Firefly luciferase gene under the control of estrogen response elements (ERE). Briefly, the cells were maintained in phenol-red-free Eagle’s minimum essential medium (MEM), supplemented with 10% dextran charcoal-stripped fetal bovine serum and 60 mg/L kanamycin, at 37°C under 5% CO_2_ atmosphere. The test medium was the same as the culture medium, but without kanamycin.

#### 2.2.3 Cytotoxicity

Cytotoxicity of the mycotoxins in the hERα-HeLa9903 cell model was assessed by using the cell proliferation reagent WST-1 according to manufacturer’s instructions (Roche, Manheim, Germany). hERα-HeLa9903 cells were plated at 1×10^4^ cells/well, 100 μL/well on a 96-well clear-bottomed plate for 3h prior to treatment. Stock solutions of mycotoxins were made in DMSO and diluted in cell culture medium. A 50 μL aliquot of each concentration was added to each well, with three replicates per condition. Plates were incubated for 24h. Then 15 μL/well (1/10 of the cell culture volume) of WST-1 was added. Cells were incubated for 4h in the dark at 37°C, 5% CO_2_. Cell viability was measured based on the conversion of tetrazolium salt to formazan by the mitochondrial dehydrogenase enzyme in metabolically active cells. Using a microplate reader (Tecan, Infinite 200 PRO, Mannedorf, Switzerland), the optical density was read at 450 nm and 690 nm. Formazan production was quantified by subtracting the signal at 690 nm from the signal obtained at 450 nm.

#### 2.2.4 ER STTA

For ER transactivation assays, hERα-HeLa9903 cells were plated at 1×10^4^ cells/well, 100 μL/well on a 96-well clear-bottomed plate for 3h prior to treatment. Stock solutions of mycotoxins, E2, OHT were made in DMSO, and diluted in cell culture medium. A 50 μL aliquot of each concentration was added to each well, with three replicates per condition. Plates were incubated for 24h. Mycotoxins were added alone (agonist assay) or in the presence of 50 pM E2 (antagonist assay). Cells were washed once with PBS 1X before adding Cell Culture Lysis reagent (E1531, Promega) and One-Glo Luciferase (E6120, Promega). Luminescence was read on a luminometer (Synergy HTX, Biotek, Agilent, Les Ulis, France).

#### 2.2.5 ER STTA Data analysis

Statistical differences between treated and solvent-control samples were assessed by one-way analysis of variance (ANOVA) and post-hoc Dunnet’s test using the GraphPad Prism software (v11.0.0; GraphPad Software, LLC, Boston, MA, USA) and setting significance at p < 0.05. Excel spreadsheets available from the OECD website were used for calculations of PC_10_ and PC_50_ (in the agonist assay, the concentration of a test substance at which the response is 10% or 50% respectively of the response induced by the positive control E2, 1 nM); as well as for calculations of IC_30_ and IC_50_ (in the antagonist assay the concentration that reduced by 30% and 50% the signal of estradiol 50 pM alone respectively). According to the OECD guideline, the endocrine effect was determined from at least two independent experiments producing similar results.

### 2.3 Androgen receptor stably transfected transactivation assay (AR STTA)

#### 2.3.1 Chemicals and reagents

Dihydrotestosterone (DHT, CAS 521-18-6, purity ≥ 95%) and mestanolone (CAS 521-11-9) were obtained from LGC (Guildford, UK). Hydroxyflutamide (HF, CAS 52806-53-8, purity ≥ 98%) and Di(2-ethylhexyl)phthalate (DEHP, CAS 117-81-7, purity ≥ 99.5%), were purchased from Sigma Aldrich (Darmstadt, Germany). Bisphenol A (BPA, CAS 80-05-7, purity > 98%) was obtained from Chiron AS (Trondheim, Norway). Cycloheximide (CAS 66-81-9, purity ≥ 95%) was from ThermoFisher Scientific (Waltham, MA, USA). Phenol-red-free DMEM/F-12 (Cat. No. D6434, with HEPES, with sodium bicarbonate, with sodium pyruvate, without L-Glutamine, without phenol red), L-Glutamine (Cat. No. G7513, 200 mM), phosphate buffered saline (Cat. No. P4474), MEM non-essential amino acid (MEM NEAA, 100x, Cat. No. M7145) and trypsin/EDTA solutions (Cat. No. T4174, 10x) were obtained from Sigma Aldrich (Darmstadt, Germany). DMSO (Cat. No. AC327182500, purity 99.8%), fetal bovine serum (FBS, Cat. No. A5256701) and penicillin-streptomycin (Cat. No. 15140-122, 10,000 U/mL) were obtained from ThermoFisher Scientific (Waltham, MA, USA). Zeocin (ant-zn-1, 100 mg/mL) and hygromycin (ant-hg-1, 100 mg/mL) were from InvivoGen (Toulouse France, Europe). HyClone™ Charcoal/Dextran treated fetal bovine serum (DCC-FBS, Cat. No. SH30068.02) was purchased from Cytiva (Global Life Sciences Solutions USA LLC, Marlborough, MA, United States). Dual-Glo® Luciferase Assay System (Cat. No. E2940) was purchased from Promega (Madison, USA).

#### 2.3.2 Cell culture

Androgenic/anti-androgenic activity of mycotoxins was assayed using the AR-EcoScreen™ cell line according to OECD TG 458 protocol (OECD 2023). The AR-EcoScreen cell line was purchased from the Japanese Collection of Research Bioresources Cell Bank (JCRB1328, JCRB, Osaka Japan). Briefly, cells were cultured in DMEM/F-12 medium, supplemented with 5% FBS, 0.1 mM MEM NEAA, 100 U/mL penicillin/streptomycin (P/S), 100 μg/mL hygromycin B, 200 μg/mL zeocin and 2.4 mM L-glutamine at 37 °C, in a humidified atmosphere containing 5% CO2. The cells were passaged twice weekly using a 0.25% trypsin/0.02% EDTA solution at a 1:8 ratio and maintained until passage 30.

#### *2.3.3 AR STTA* and cytotoxicity

For the AR transactivation assays, cells were seeded at a density of 1×10^5^cells/ml in 96-well white microtiter plates (Nunclon Delta-Treated, Flat-Bottom) in 100 μL phenol red-free DMEM/F-12 medium supplemented with 5% DCC-FBS, 100 units/mL P/S and 2.4 mM L-glutamine (treatment medium). Cells were incubated overnight to allow attachment. At the time of treatment, the medium was removed and replaced with freshly prepared treatment medium supplemented with *Alternaria* toxins at final concentrations ranging from 0.001 to 60 μΜ or with appropriate positive/negative controls. Working solutions were prepared by serial dilutions of the stock solution in treatment medium supplement with 0.1% (v/v) DMSO for the agonism assay or in treatment medium containing 500 pM DHT and 0.1% (v/v) DMSO for the antagonism assay. Cytotoxicity control (10 μg/ml cycloheximide) was also included in each run. Cells were incubated for 22-24h.

Both AR response (Firefly luciferase activity) and cell viability (*Renilla* luciferase activity) were determined sequentially using the Dual-Glo Luciferase Assay system (Promega, Madison, WI, USA) in white, opaque, 96-well plates. Luminescence was measured using the Varioskan™ LUX Multimode Microplate Reader (Thermo Scientific) employing default instrument parameters. At least two valid independent experimental runs were performed for each test item, according to OECD TG 458 protocol (OECD 2023).

*Alternaria* toxins were tested at seven concentrations together with solvent control, positive controls and complete concentration-response curves of the reference items. All assay acceptability criteria were met and the reference items were correctly classified as positive (DHT, mestanolone for agonism and HF, BPA for antagonism) or negative (DEHP) according to OECD TG 458 (OECD 2023). In line with the OECD TG 458, co-incubation with hydroxyflutamide (HF) was performed to confirm involvement of the AR and exclude glucocorticoid receptor (GR) cross talk.

#### 2.3.4 AR STTA data analysis

For AR response assessment, recorded Firefly luminescence signals were corrected by subtracting the mean solvent control value and then expressed as percentages of respective positive control values (10 nM DHT for agonism and 0.5 nM DHT for antagonism), set at 100%. For the assessment of cell viability, recorded *Renilla* luminescence signals were corrected by subtracting the mean positive control for cytotoxicity (10 μg/ml cycloheximide) and then expressed as percentages of mean solvent control value, set at 100%. Concentrations that reduced *Renilla* luciferase activity by 20% or more were regarded as cytotoxic, according to OECD TG 458 (OECD 2023).

The effective concentrations of positive controls (PC_10_, PC_50_ values for agonism and IC_30_, IC_50_ for antagonism) and the concentrations resulting in 50% reduction in viability (IC_50_) were calculated using drc v3.0 in R v4.5.2, applying the three parameters log-logistic curve fitting, following OECD TG 458 instructions for data analysis. Statistical analysis was performed using Student’s t-test, two-tailed distribution, assuming two-sample unequal variance.

### 2.4. H295R Steroidogenesis assay

#### 2.4.1 Chemicals and reagents

Prochloraz (PRO, CAS 67747-09-5, purity ≥ 98%), forskolin (FOR, CAS 66575-29-9, purity ≥ 95%) and bisphenol A (BPA, CAS 80-05-7, purity > 98%) were purchased from Sigma Aldrich (Darmstadt, Germany). Atrazine (ATZ, CAS 1912-24-9, purity ≥ 98%) was from Sigma Aldrich, aminoglutethimide (AMG, AS 125-84-8, purity ≥ 99%) was from Santa Cruz (Heidelberg, Germany). Phenol red-free DMEM/F12 medium containing L-glutamine and HEPES (Cat. No. 11039021) was purchased by Gibco/ThermoFisher Scientific (Dreieich, Germany), ITS-Premix (Cat. No. 11593560), and Nu-serum IV (Cat. No. 11563600) from Fisher Scientific (Schwerte, Germany), penicillin/streptomycin solution (100 U/mL) were from Capricorn Scientific (Ebsdorfergrund, Germany), DMSO (Cat. No. 1029521000) was from Merck (Darmstadt, Germany), ELISA kits (Cat No. E2: DE2693; T: DE1559) were from Demeditec Diagnostics (Kiel, Germany), 3-(4,5-dimethylthiazol-2-yl)-2,5-diphenyltetrazolium bromide (MTT) (Cat. No. M2128), and Triton X-100 (Cat. No. T8787) were from Merck.

#### 2.4.2 Cell culture

H295R cells (ATCC, Manassas, VA, USA) were maintained in phenol red-free DMEM/F12 medium containing L-glutamine and HEPES supplemented with 1% (v/v) ITS-Premix, 2.5% (v/v) Nu-serum IV, 100 U/mL penicillin, and 100 μg/mL streptomycin in a humidified atmosphere at 5% CO_2_ at 37°C. The cells were cultured according to the standardized protocol approved by the OECD (OECD 2025b) (OECD, 2025). The medium changed every 2 to 3 days and cells were splitted once per week in 1:3 ratio when 80-90% confluence was reached. The cells were used for testing from passage 5 to 10.

#### 2.4.2 Cytotoxicity

Prior to the steroidogenesis assay, range finding tests for cytotoxicity were performed. H295R cells were seeded in 96-well cell culture plates at a density of 60.000 cells/well/100 µL. After 24h, the culture medium was removed and fresh medium containing various concentrations of the selected *Alternaria* toxins were added. DMSO (1% v/v) served as solvent control and 0.01% Triton X-100 as positive control for cytotoxicity. After 48h of treatment, the cellular metabolic activity as an indirect measure of cellular viability was assessed using the MTT assay as previously described (Behr et al. 2018) (Behr et al., 2018).

#### 2.4.3 Steroidogenesis assay

The H295R cells (60000 cells/well/100 µL) were seeded in 96-well plates, incubated for 24h, and subsequently treated with seven concentrations of the *Alternaria* toxins in triplicates for 48h. The medium was collected in 0.2 mL tubes and stored at -80°C until hormone analysis. A quality control plate was included in each run containing blanks (only culture medium), solvent control (0.1% v/v DMSO), FOR (1 µM, 10 µM) and PRO (0.1 µM, 1 µM) as a known inducer and inhibitor for E2 and T production. E2 and T levels were measured using ELISA kits (Demeditec Diagnostics) according to the manufacturer. Three individual experiments were conducted.

Potential interference by matrices and chemicals were excluded by performing the respective tests according to the guideline. To demonstrate the performance of the assay, a proficiency test was performed with various concentrations of FOR, PRO, ATZ, AMG, and BPA.

#### 2.4.4. Steroidogenesis assay data analysis

Results were considered positive when there was a statistically significant fold change (fc) of ≥ 1.5 in at least two adjacent concentrations compared to solvent control. Statistical differences between treatment and control were only determined if fc ≥ 1.5 using ANOVA with a Dunnett’s post hoc test. IC_50_ values were calculated by fitting the experimental data to the 4-parameter logistic (4PL) equation using GraphPad prism 10.1.2.

### 2.5 E-Screen assay

#### 2.5.1 Chemicals and reagents

Fulvestrant (ICI 182,780, CAS 129453-61-8, purity ≥ 98%), E2 (CAS 50-28-2, purity ≥ 98%), L-Glutamine (Cat. No. G7513, 200 mM), phosphate buffered saline (Cat. No. P4474), trypsin/EDTA solution (Cat. No. T4174, 10X) and MTT (Cat. No. M5655) were obtained from Sigma Aldrich (Darmstadt, Germany). RPMI 1640 medium (Cat. No. 21875034, with glutamine, with phenol red, with sodium bicarbonate, without HEPES, without sodium pyruvate), RPMI 1640 medium (Cat. No. 11835063, with glutamine, with sodium bicarbonate, without HEPES, without phenol red, without sodium pyruvate), DMSO (Cat. No. AC327182500, purity 99.8%), fetal bovine serum (FBS, Cat. No. A5256701), penicillin-streptomycin solution (Cat. No. 15140-122, 10000 U/mL) and CyQUANT® Direct Cell Proliferation Assay (Cat. No. C35011) were obtained from Thermo Fisher Scientific (Waltham, MA, USA) and DCC-FBS (Cat. No. SH30068.02) from Cytiva (Global Life Sciences Solutions USA LLC, Marlborough, MA, United States).

#### 2.5.2 Cell culture

MCF-7 cells were obtained from ATCC and maintained in T-75 culture flasks at 37°C in a humidified incubator containing 5% CO_2_ and 95% air. The cells were routinely cultured RPMI 1640 medium supplemented with 10% FBS, 1% penicillin and streptomycin solution, according to standard cell culture procedures. Prior to treatment with the test compounds, cells were cultured in phenol red-free medium containing 5% DCC-FBS.

#### 2.5.3 E-Screen assay

Cell proliferation was determined using the E-Screen assay (Soto et al. 1995). MCF-7 cells were seeded in 96-well plates at a density of 5×10^4^ cells/mL, 100μL/well. After 24h, the cells were treated with different concentrations of the test compounds (0.001 μM to 100 μΜ). E2 (0.001 μM) was used as positive control. Co-incubation of mycotoxins with E2 (0.001 μM) was also performed. After three days of treatment, the medium was replaced with fresh RPMI containing the test compounds. Cell proliferation was assessed after six days of treatment using the CyQUANT Direct Cell Proliferation Assay, which measures cellular DNA content using a cell-permeant nucleic acid dye in combination with a background suppression reagent, providing a fluorescence-based proxy for cell number. The assay was performed according to the manufacturer’s instructions. Briefly, equal volumes of detection reagent were added directly to the cell culture medium, and the plates were incubated for 60 min at 37°C before fluorescence measurement. Fluorescence was measured using excitation and emission wavelengths of 480 nm and 535 nm, respectively, with bottom-read detection in the VarioScan Lux microplate reader (Thermo Fisher Scientific Waltham, MA, USA). Fluorescence intensity was proportional to the number of viable cells and was normalized to untreated (solvent) control samples, which were considered 100% viable. Statistical analysis was performed using Student’s t-test, two-tailed distribution, assuming two-sample unequal variance.

## 3. Results

### 3.1 ER STTA

In the agonistic part of the estrogen receptor transcriptional activation assay using hERα-HeLa9903 cells, TEN, ALT, TeA and ATX-I did not induce ER-mediated activation as luminescence signals did not reach 10% of that of 1 nM E2 alone, as required by the OECD TG 455 positivity criteria (**Figure 1**). On the other hand, AOH and AME induced a significant activation of the estrogen receptor (PC_10_: 1.1 - 1.3 µM and PC_50_: 3.9 - 4.6 µM; PC_10_: 1.4 - 1.5 µM and PC_50_: 5.2 - 8.5 µM respectively) at concentrations inducing less than 20% cytotoxicity. The calculated IC_50_ for cytotoxicity were higher than 30 µM for TeA, TEN, ALT, AOH and AME and 30.3 ± 4.0 µM for ATX-I.

**Figure 1:**
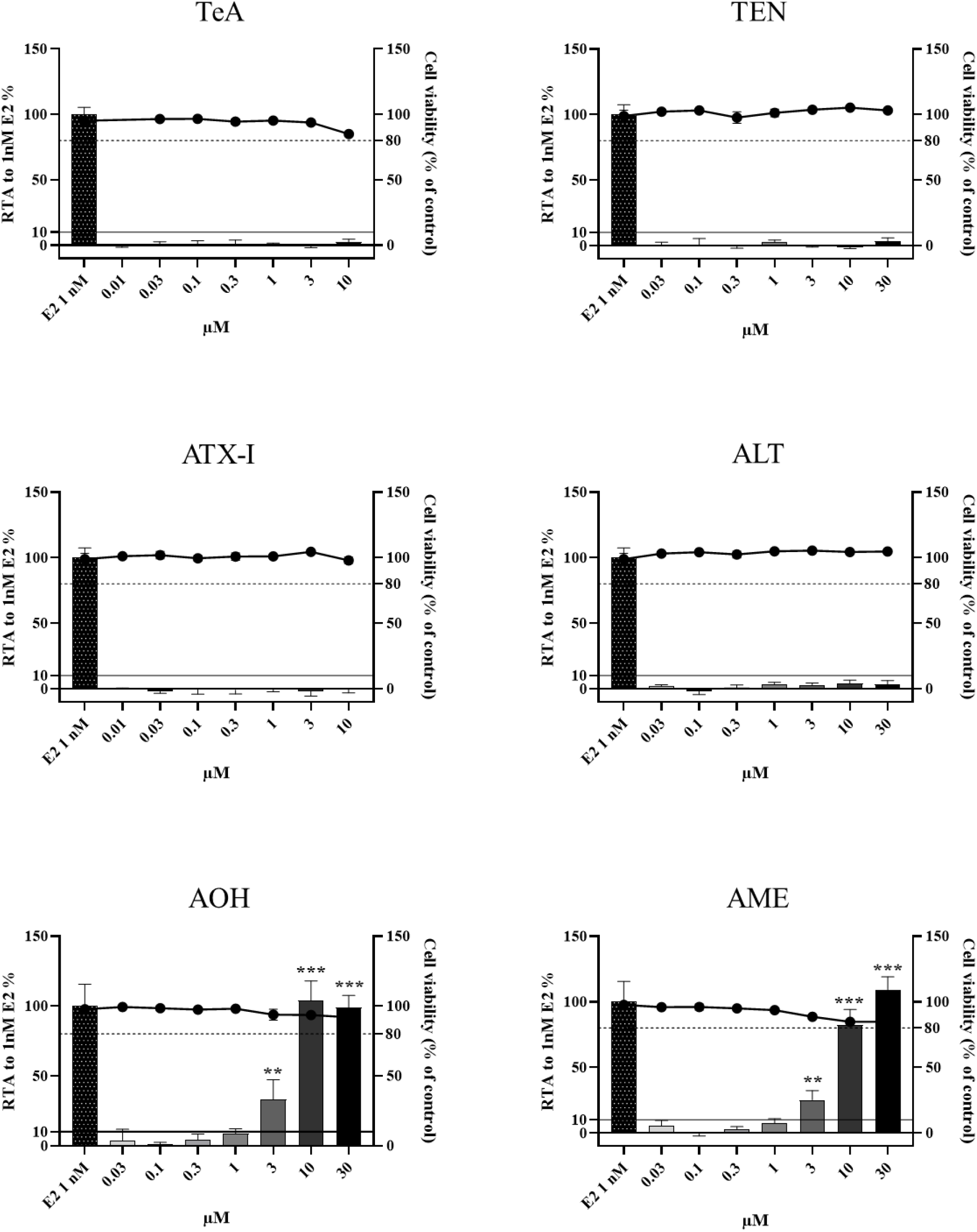
Agonistic properties (bars) and cytotoxicity (solid line) of TeA, TEN, ATX-1, ALT, AOH and AME on the estrogen-responsive cell line hERα-HeLa9903 cells, following treatment at the indicated concentrations of mycotoxins using the One-Glo Luciferase assay. Bars represent mean ± SD % Relative transcriptional activity (RTA%) normalized to the E2 positive control (1 nM). Solid line represents mean ± SD % cell viability normalized to solvent control (DMSO 0.1%). The asterisks indicate the level of significance: ** p<0.01, *** p < 0.001.

In the antagonist assay, none of the mycotoxins exhibited antagonistic activity, except for ATX-I which was shown to induce a statistically significant inhibition (IC_30_: 0.27 - 0.37 µM). However, the response remained below 50% compared to E2 (50 pM) alone (**Figure 2**). The positive control OHT (1 µM) induced a strong inhibition of the transcriptional activity of the ER, reducing the relative transcriptional activity of E2 (50 pM) by at least 60% in good agreement with the acceptability criteria of the assay. The well-known ER antagonist tamoxifen induced a significant inhibition of the relative transcriptional activity of ER (IC_30_: 0.21 - 0.54 µM; IC_50_: 0.55 - 1.32 µM). Interestingly, AOH and AME, from the concentration of 3µM, induced an enhanced signal compared to E2 alone (50 pM). TeA at the highest concentration tested also increased the signal above that of E2 control alone, although this compound was not identified as an agonist when tested alone.

**Figure 2:**
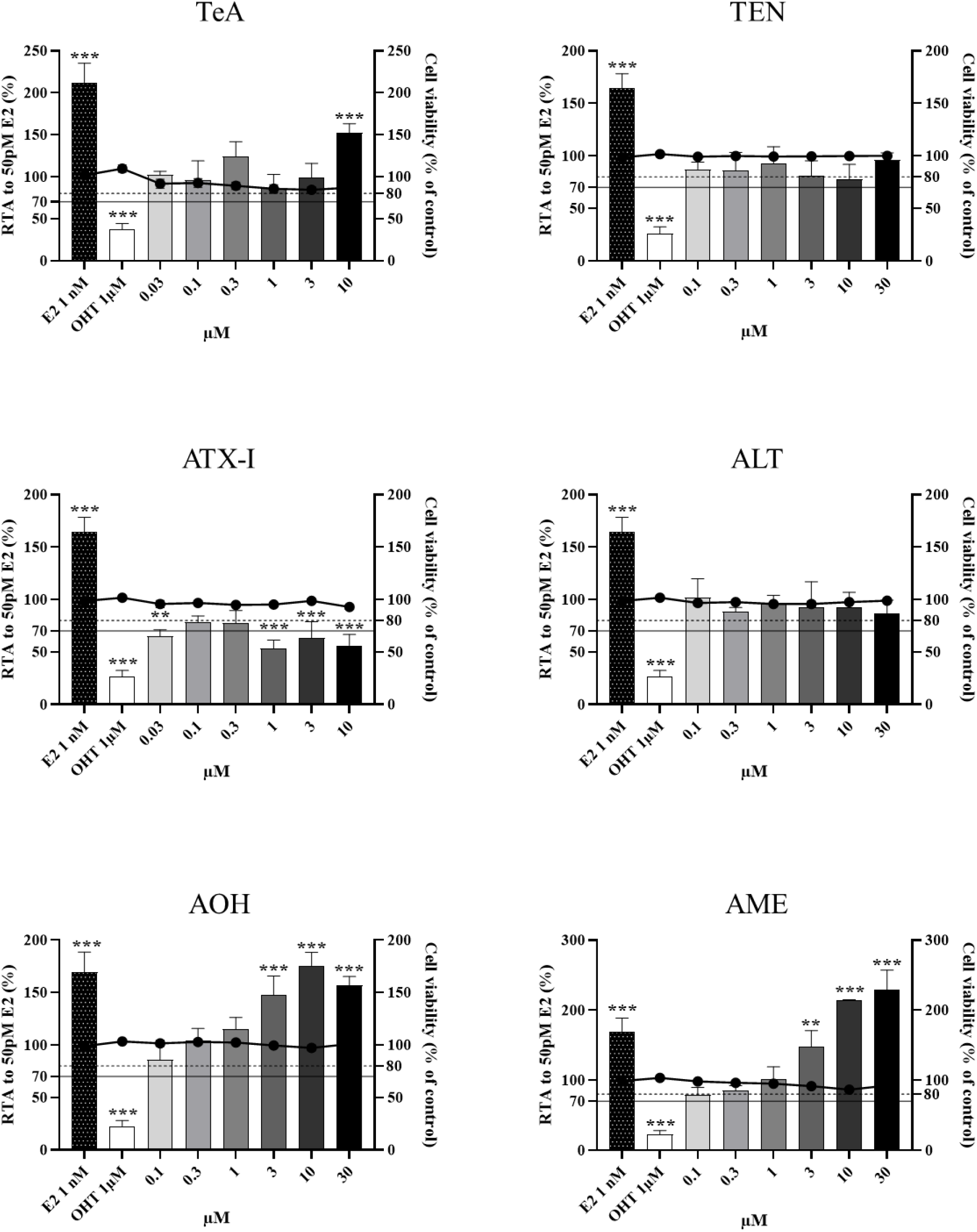
Antagonistic properties (bars) and cytotoxicity (solid line) of TeA, TEN, ATX-I, ALT, AOH and AME on the estrogen-responsive cell line hERα-HeLa9903 cells, following exposure at the indicated concentrations of mycotoxins using the One-Glo Luciferase assay. Bars represent mean ± SD % Relative transcriptional activity (RTA%) normalized to the E2 control (50 pM). Solid line represents mean ± SD % cell viability normalized to solvent control (DMSO 0.1%). OHT: 4-Hydroxytamoxifen. Asterisks indicate the level of significance: ** p<0.01, *** p < 0.001.

### 3.2 AR STTA

TeA, ALT, TEN and ATX-I were judged negative for AR agonistic effects as no increases were observed at non cytotoxic concentrations (**Figure 3**). Across three independent experimental runs, AOH and AME induced statistically significant increases in luciferase activity exceeding 10% of the positive control at high concentrations, while cytotoxicity remained below the 20% threshold (**Figure 3, Supplementary Information SI-1**). Based on the OECD TG 458 decision criteria, both AOH and AME were considered positive for AR-mediated agonistic activity. The calculated PC_10_ values ranged between 3.53 - 12.5 μΜ for AOH and 6.87 - 19.7 μΜ for AME suggesting a weak response. Marked cytotoxicity was observed at concentrations above 10 μΜ for ΑΤΧ-Ι and above 30 μΜ for AOH and AME. IC_50_ values for cytotoxicity could be estimated only for ATX-I at 19.04 ± 5.30 μM and at 46.93 ± 7.69 μM for AOH. No cytotoxicity was observed for ALT and TEN at concentrations up to the maximum solubility (24 and 34 μΜ respectively).

**Figure 3:**
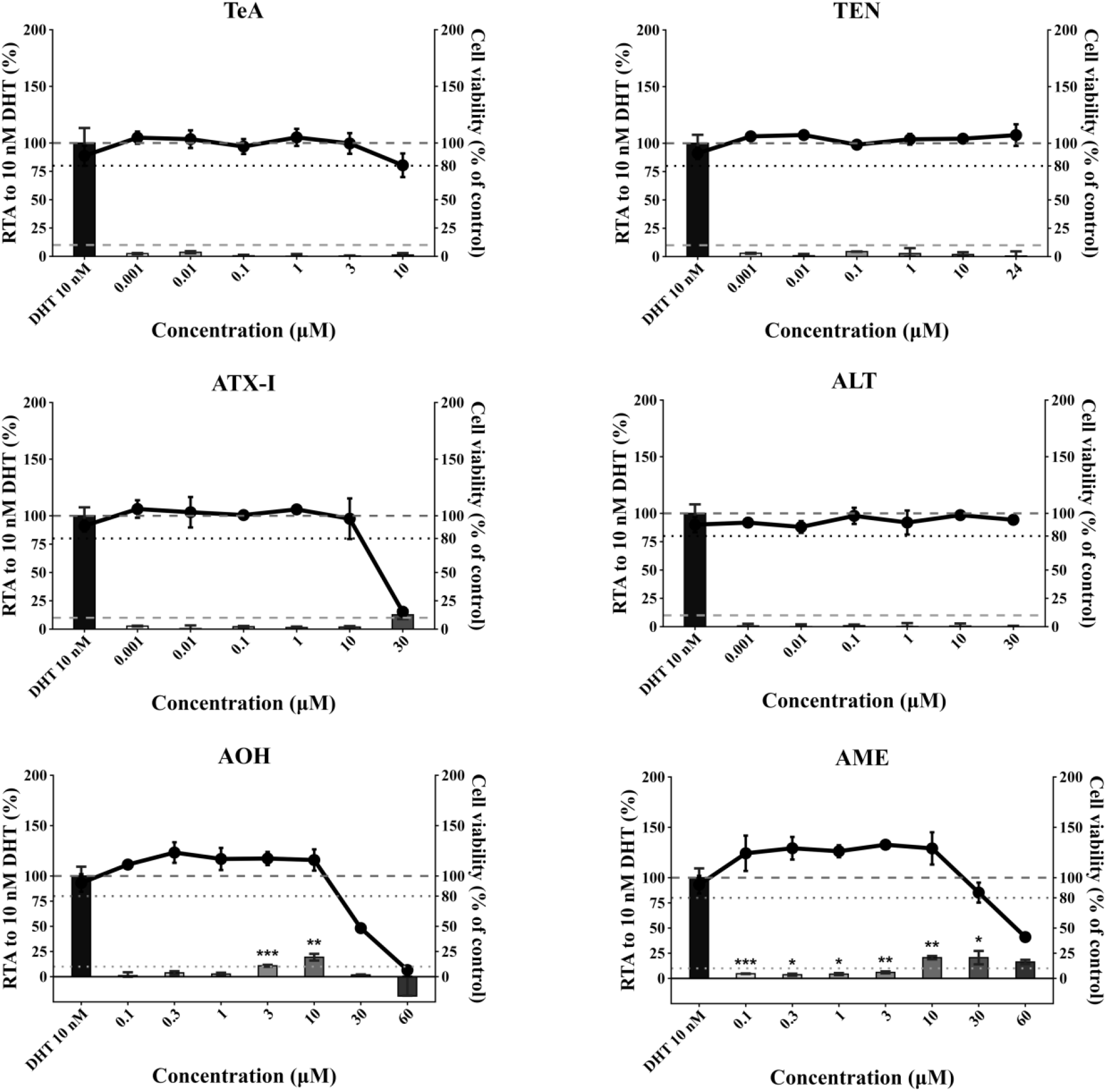
Agonistic (bars) and cytotoxicity (solid line) of TeA, TEN, ATX-I, ALT, AOH and AME in the AR-EcoScreen cell line, following exposure at the indicated concentrations. Bars represent mean ± SD % Relative Transcriptional Activity (RTA) of at least two independent experiments (TeA, TEN, ATX-I, ALT) with consistent results or mean ± SD % RTA of one representative experiment (AOH, AME) normalized to the reference compound dihydrotestosterone (DHT, 10 nM). Dots represent mean ± SD % cell viability normalized to solvent control (0.1% DMSO). AR response (Firefly luciferase activity) and cell viability (*Renilla* luciferase activity) were determined using the Dual-Glo Luciferase Assay. Asterisks indicate the level of significance: *p<0.5, ** p<0.01, *** p < 0.001 (compared to solvent control)

As minimal GR-mediated crosstalk may occur in the AR-EcoScreen cell line, co-incubation with the specific AR antagonist hydroxyflutamide (HF) was performed for compounds judged as positive to determine whether the weak induced responses could be blocked. As presented in **Figure 4**, AOH and AME alone induced luciferase activity above 10% of the positive control from the dose of 10 μΜ. For the AR agonist DHT, a marked decrease in luciferase activity was observed in the presence of 1 μΜ HF. However, only a mild decrease in respective activity was observed for AOH and AME co-incubated with 1 μΜ HF. In addition, AME was still considered positive in line with OECD criteria (above 10% of 0.5 nM DHT response) suggesting that the observed minimal activation could be, at least in part, mediated by GR. This was further supported by the “true” AR-responses that were calculated according to OECD TG 458 in the presence of 1 μΜ HF and were below the 10% threshold for positivity (**Supplementary Information SI-2**).

**Figure 4:**
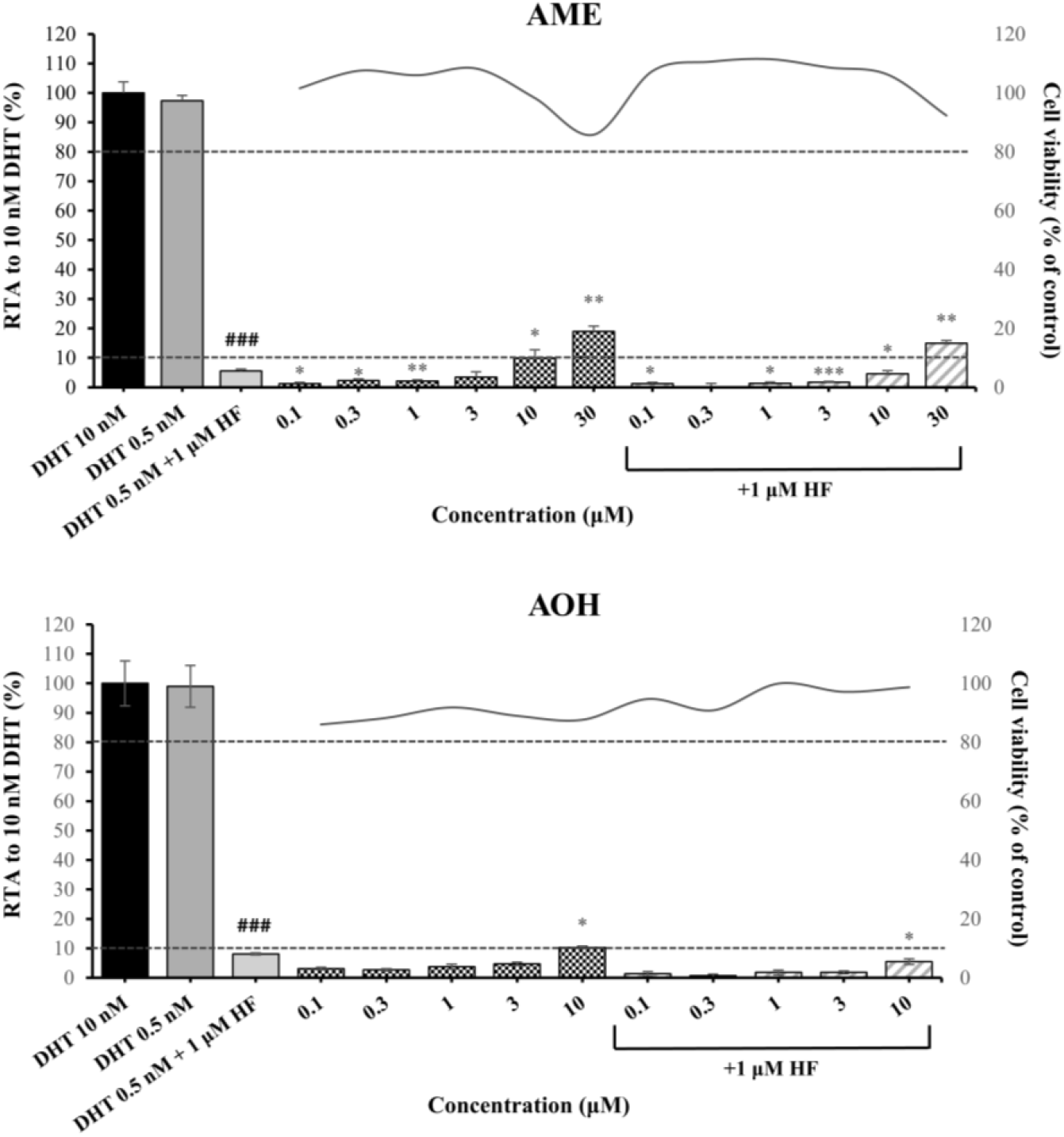
Agonistic properties (bars) and cytotoxicity (solid line) of AOH and AME in the AR-EcoScreen cell line in the presence of the AR antagonist HF. Bars represent mean ± SD % Relative Transcriptional Activity (RTA) of one representative experiment normalized to the reference compound dihydrotestosterone (DHT, 10 nM). AR response (Firefly luciferase activity) and cell viability (*Renilla* luciferase activity) were determined using the Dual-Glo Luciferase Assay. HF: Hydroxyflutamide. Asterisks indicate the level of significance: *p<0.5, ** p<0.01, *** p < 0.001 (compared to solvent control); ^###^ p < 0.001 (compared to DHT)

Regarding AR antagonistic effects, decreases in AR-mediated transcriptional activation in the presence of DHT were observed for AOH, AME, ATX-I, and TeA at concentrations that also induced cytotoxicity. The 50% decrease in DHT-induced transcriptional activation observed for AME at 30 μM is considered questionable, as cytotoxicity at this concentration (18.6%) was close to the 20% threshold set by OECD TG 458. Moreover, marked cytotoxicity of up to 40% was observed at the next tested concentration of 60 μM. Thus, all *Alternaria* compounds were judged as negative for AR-mediated antagonistic effects (**Figure 5**).

**Figure 5:**
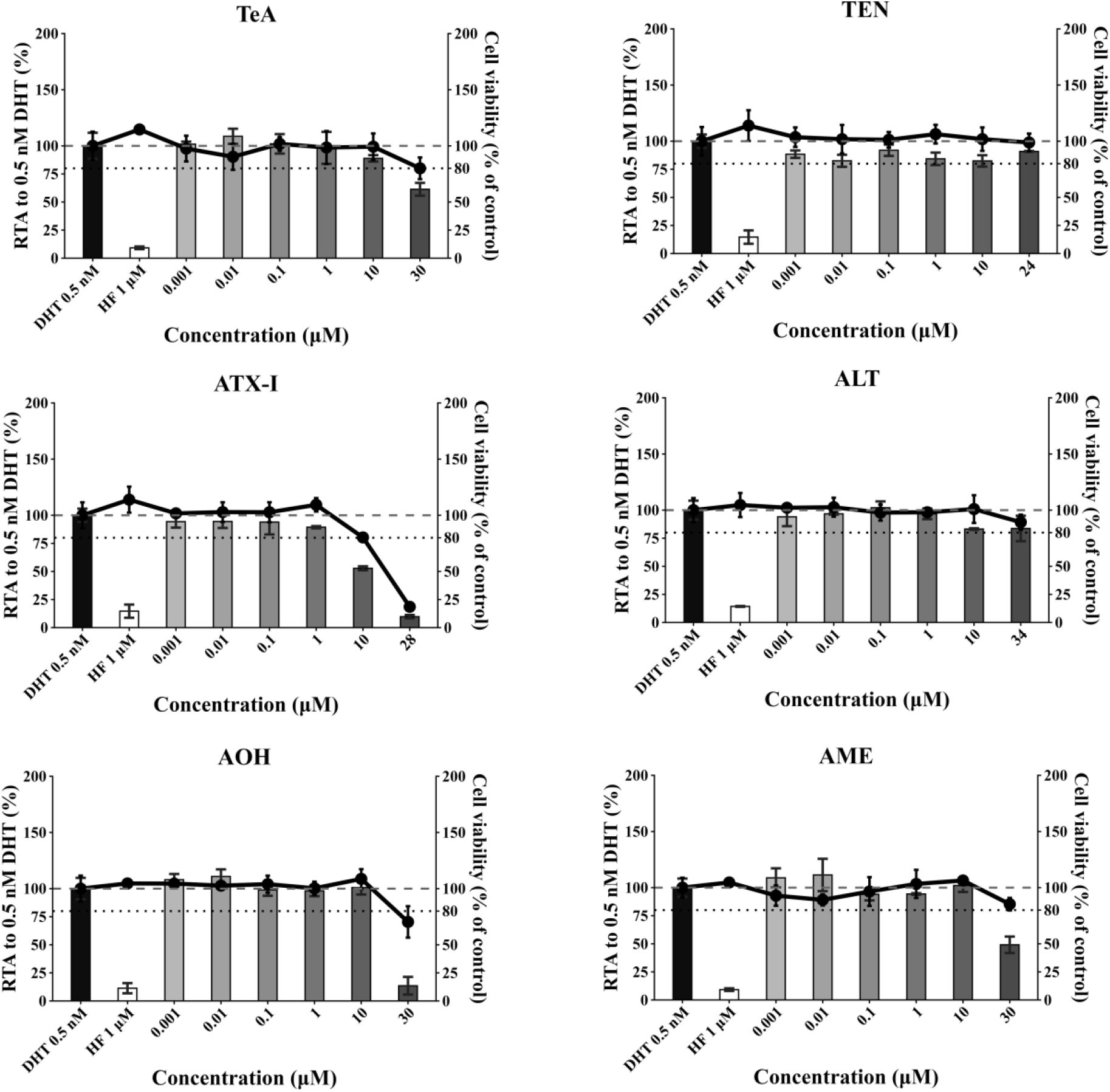
Antagonistic (bars) and cytotoxicity (solid line) of TeA, TEN, ATX-I, ALT, AOH and AME in the AR-EcoScreen cell line, following exposure at the indicated concentrations. Bars represent mean ± SD % Relative Transcriptional Activity (RTA) of two independent experiments with consistent results normalized to the reference compound dihydrotestosterone (DHT, 0.5 nM). Dots represent mean ± SD % cell viability normalized to solvent control (0.1% DMSO). AR response (Firefly luciferase activity) and cell viability (*Renilla* luciferase activity) were determined using the Dual-Glo Luciferase Assay. HF: Hydroxyflutamide

Suitability of the assay system was demonstrated using the OECD TG 458 reference/profficiency substances DHT, mestanolone, and DEHP for the agonistic part and HF, BPA and DEHP for the antagonistic part (**Supplementary Information SI-3**).

### 3.3 Steroidogenesis assay

#### Cytotoxicity of *Alternaria* toxins in H295R cells

For range finding prior to the steroidogenesis assay, the cytotoxicity of the selected *Alternaria* toxins was assessed using the MTT assay as an indirect measure of the cell viability based on cellular metabolic activity (**Figure 6**). TeA and ATX-I showed a concentration-dependent decrease in the metabolic activity of the H295R cells, whereas TEN and ALT did not affect metabolic activity at any of the tested concentrations. For AOH and AME, precipitate formation was observed at the highest tested concentration (150 µM and 50 µM, respectively). The calculated IC_50_ values were 114.2 µM for ATX-I and 4873 µM for TeA, the latter being extrapolated from the concentration-response curve. For AME, an IC_50_ value of 32.48 µM was obtained. However, this value should be interpreted with caution, due to precipitate formation. No IC_50_ values could be determined for TEN, ALT, or AOH within the tested range.

**Figure 6:**
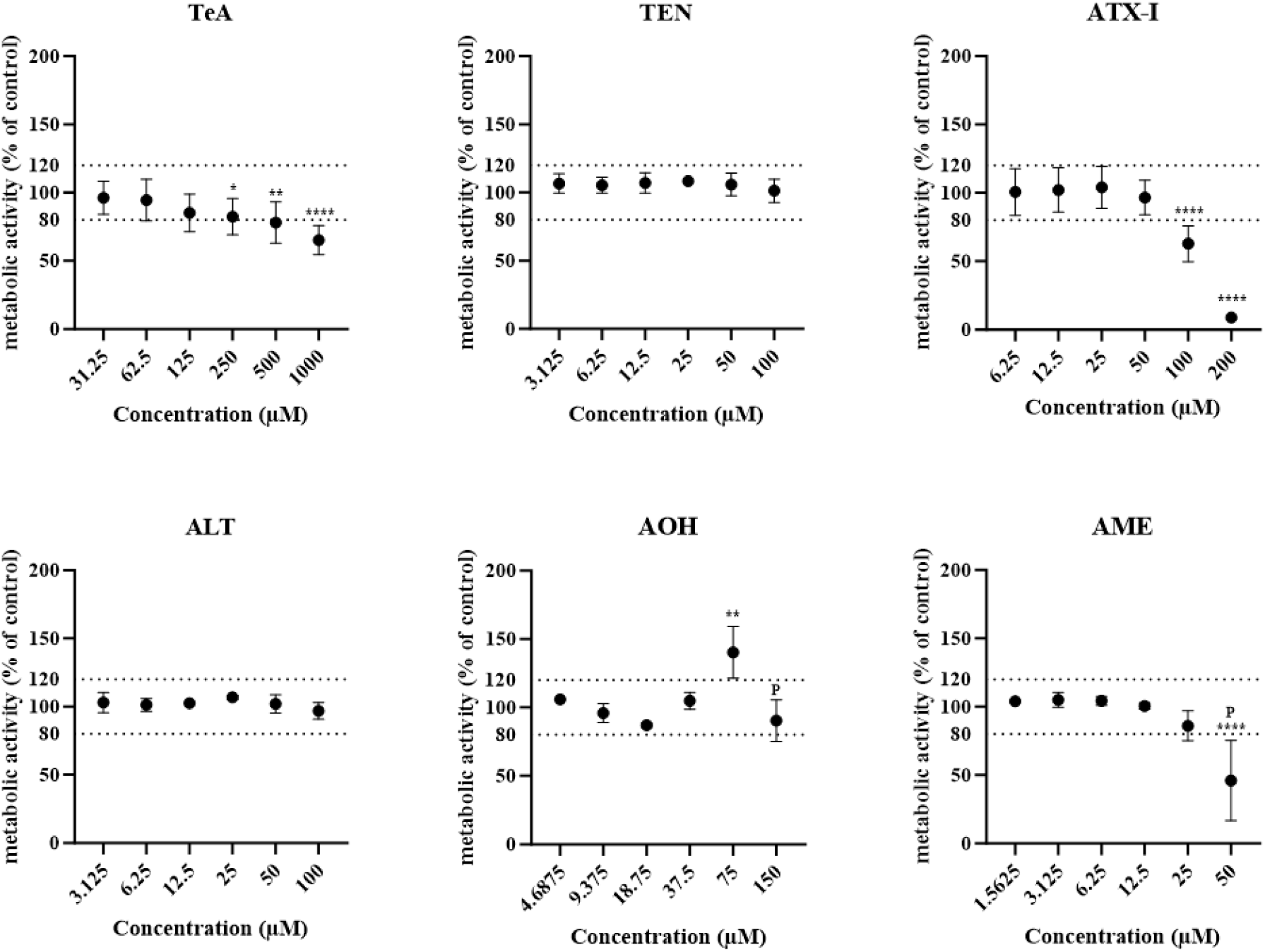
Cytotoxicity of TeA, TEN, ATX-I, ALT, AOH and AME on H295R cells after 48h treatment using the MTT assay. Dots represent the mean ± SD metabolic activity normalized to the solvent control (DMSO 0.1%) in percent. P: Precipitates. Asterisks indicate the level of significance: *p<0.05, **p<0.01, ****p<0.0001.

#### Impact of *Alternaria* toxins on E2 and T levels in H295R cells

To determine the impact of the selected *Alternaria* toxins on E2 and T production, the steroidogenesis assay was performed in H295R cells according to OECD TG 456. None of the tested *Alternaria* toxins induced biologically relevant alterations in hormone production, as no changes exceeded the OECD threshold of 1.5-fold increase or decrease were observed for either E2 (**Figure 7**) or T levels (**Figure 8**). Decision matrix according to the OECD criteria is provided in **Supplementary Information SI-4**.

**Figure 7:**
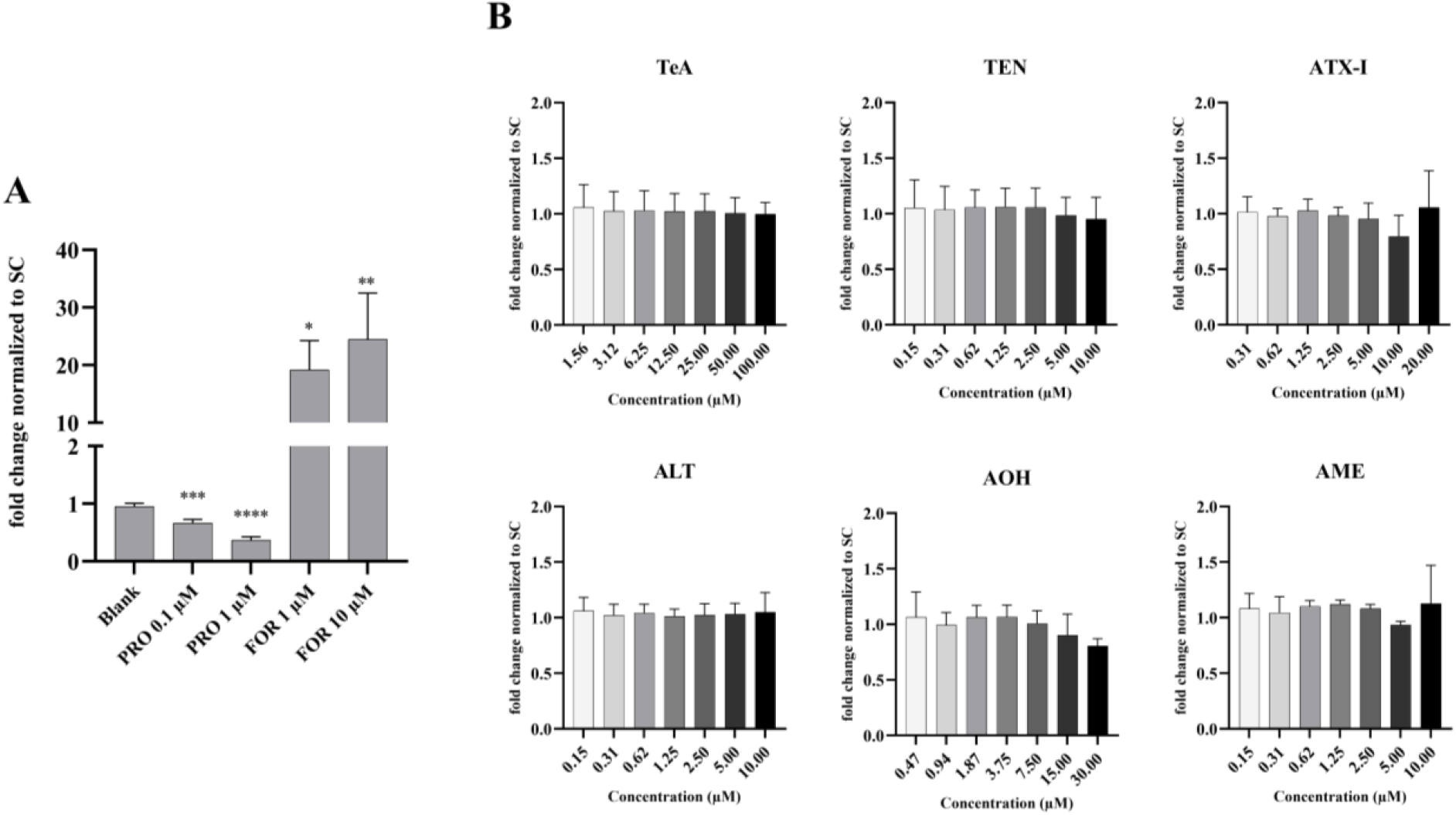
Impact of TeA, TEN, ATX-I, ALT, AOH, and AME on the hormone levels of estradiol (E2) determined via ELISA in H295R cells after 48h treatment. Bars represent the mean + SD E2 levels normalized to the solvent control out of 3 biological replicates. **A**: E2 levels of the controls with blank (cells with only medium), prochloraz (PRO) and forskolin (FOR), **B**: E2 levels after treatment with the selected *Alternaria* toxins in various concentrations. Asterisks indicate the level of significance: *p<0.05, **p<0.01, ***p<0.001, ****p<0.0001.

**Figure 8:**
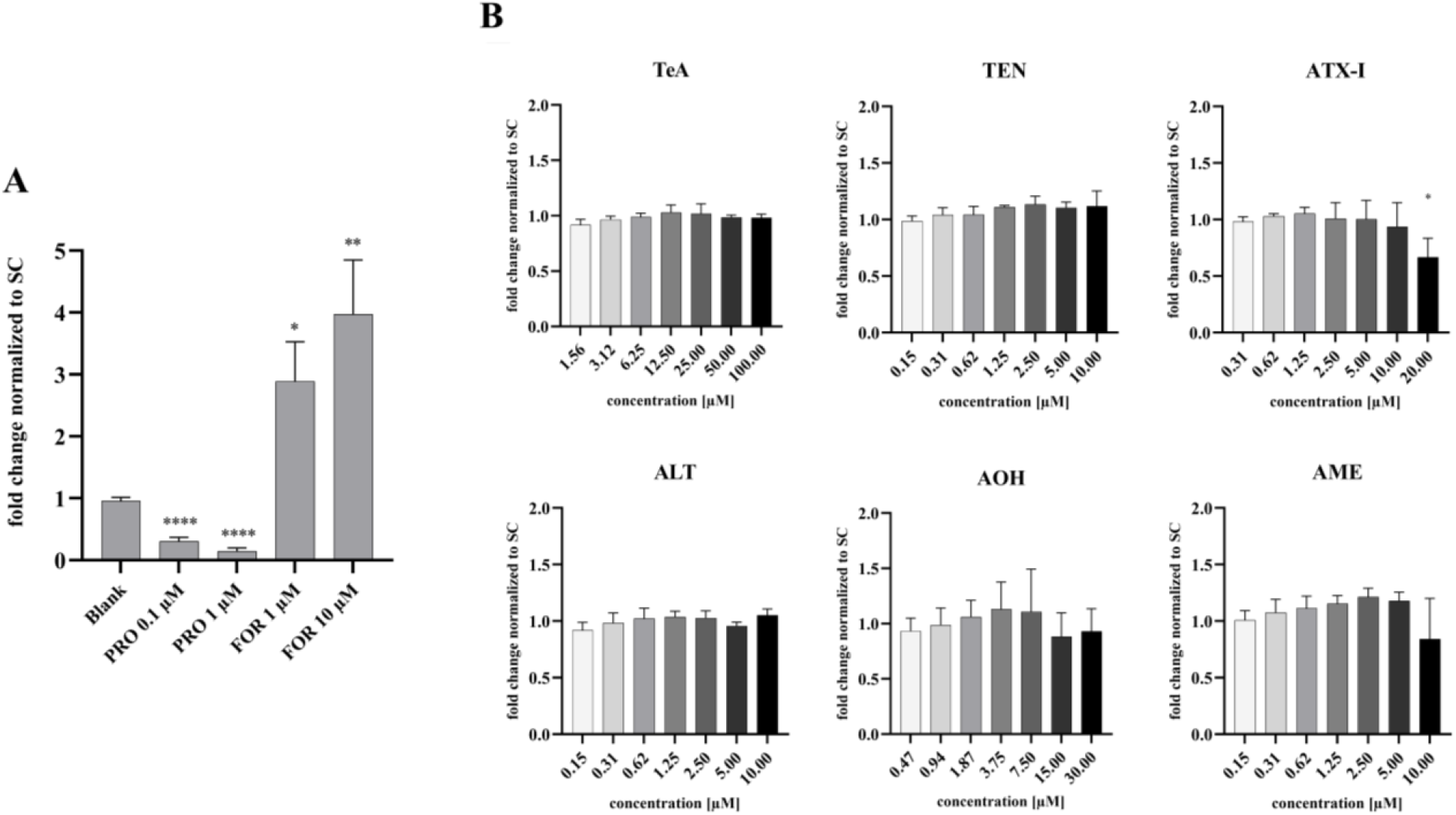
Impact of TeA, TEN, ATX-I, ALT, AOH, and AME on the hormone levels of testosterone (T) determined via ELISA in H295R cells after 48h treatment. Bars represent the mean + SD T levels normalized to the solvent control out of 3 biological replicates. **A**: T levels of the controls with blank (cells with only medium), prochloraz (PRO) and forskolin (FOR), **B**: T levels after treatment with the selected *Alternaria* toxins in various concentrations. Asterisks indicate the level of significance: *p<0.05, **p<0.01, ****p<0.0001.

The assay performance was confirmed by the expected responses of the controls. Basal hormone production in blank cultures ranged from approximately 80 to 190 pg/mL for E2 and from 5 to 8 ng/mL for T. FOR increased E2 production up to 19-fold (1 µM) and 24-fold (10 µM), while T levels increased up 2.9-fold and 4.0-fold, respectively. In contrast, PRO reduced E2 production to 0.6-fold and 0.4-fold and T production to 0.3-fold and 0.1-fold of the SC at 0.1 µM and 1 µM, respectively. In addition, assay proficiency was demonstrated using the OECD TG 456 reference substances FOR, PRO, AMG, BPA, and ATZ. All reference substances elicited the expected concentration-responses (**Supplementary Information SI-5**), confirming the suitability of the assay system.

### 3.4. E-Screen assay

All compounds tested except TeA caused a statistically significant increase in MCF-7 cell proliferation, which was confined to the highest non cytotoxic concentration only (**Figure 9**). No effect was observed at lower concentrations for any of the compounds tested. Marked cytotoxicity was observed for ATX-I, AOH and AME at the highest concentration of 100 μΜ.

**Figure 9:**
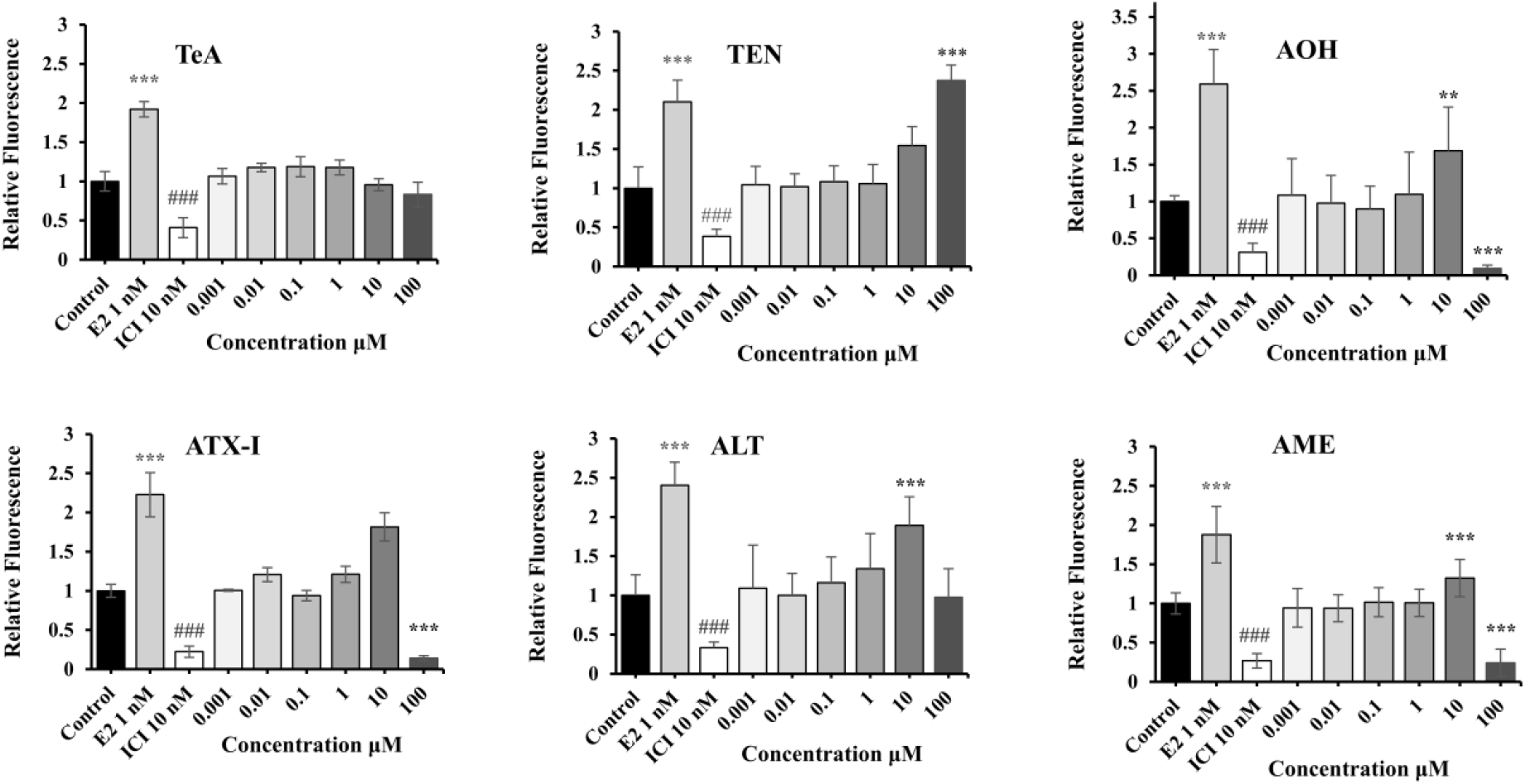
Effects of TeA, TEN, AOH, ATX-I, ALT and AME on MCF-7 cell proliferation after 6 days of treatment, assessed using the CyQUANT Direct Cell Proliferation Assay. Bars represent the mean ± SD relative fluorescence intensity (relative to the solvent control; 0.1% DMSO), used as a proxy for cell number, out of 3 biological replicates. Asterisks indicate the level of significance: *p<0.05, **p<0.01, ***p<0.001 (compared to solvent control), ^###^ p < 0.001 (compared to E2 1 nM)

Statistically significant decreases in MCF-7 cell proliferation were observed for ATX-I and TeA after co-incubation with E2 at concentrations that were not cytotoxic when tested alone (**Figure 10**). No significant decreases in breast cancer cell proliferation were observed for the rest of compounds at non-cytotoxic concentrations (data not shown).

**Figure 10:**
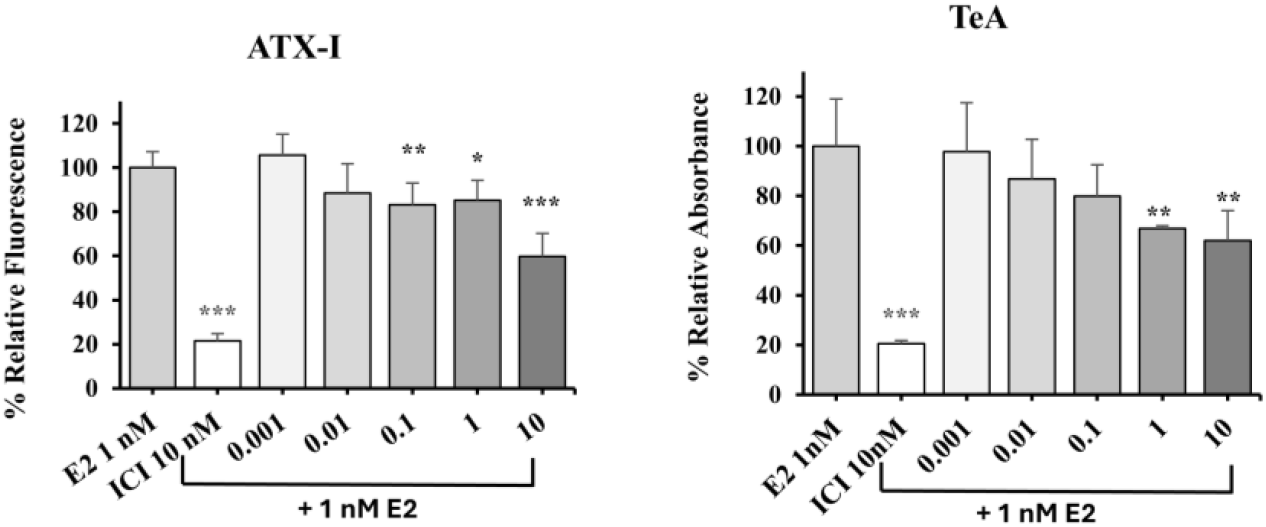
Effects of ATX-I and TeA on the induced proliferation of MCF-7 cells after co-incubation with E2 (1 nM) and 6 days of treatment, assessed using the CyQUANT Direct Cell Proliferation Assay. Bars represent the mean ± SD relative fluorescence intensity (percent relative to the solvent control; 0.1% DMSO), used as a proxy for cell number, out of 3 biological replicates. Asterisks indicate the level of significance: *p<0.05, **p<0.01, ***p<0.001 (compared to E2 1 nM)

## 4. Discussion

*Alternaria* mycotoxins are among the most widespread contaminants in food and feed worldwide, representing a potential risk to human and animal health [2]. Despite growing concern about their potential to disrupt the endocrine system, available data remain limited [6]. Following the strategy described in the EFSA/ECHA/JRC guidance for the identification of endocrine disruptors (EDs), fulfillment of the ED criteria requires the establishment of a biologically plausible link between endocrine adversity observed at the organism level and endocrine activity [16]. The evaluation of endocrine activity for the Estrogen, Androgen and Steroidogenesis (EAS)-modalities mainly relies on validated assays performed according to OECD Test Guidelines (TGs), as also described in the OECD Revised GD 150 (OECD 2018). In an effort to address the current regulatory gaps for *Alternaria* toxins, well-characterized compounds have been used to generate data on EAS-mediated activity using standardized *in vitro* assays that were conducted according to OECD guidelines.

AOH, AME, ATX-I, TEN, ALT and TeA have been assessed for possible ER-mediated (ant)-agonistic effects in the hERα-HeLa-9903 cell line in accordance with OECD TG 455. Both AOH and AME were judged positive for ER-induced agonistic activity in line with the OECD criteria with calculated PC_50_ values ranging from 3.9 to 4.6 µM for AOH and 5.2 to 8.5 µM for AME. AOH and AME also enhanced the observed induction in the presence of estradiol from the concentration of 3 µM, which is in good agreement with the agonistic effects of these toxins on the estrogen receptor. In line with these findings, AOH and AME have been reported as weak ER agonists in different test systems (Dellafiora et al. 2018; Demaegdt et al. 2016; Frizzell et al. 2013; Stypuła-Trębas et al. 2017). Comparable EC_50_ values for AOH, ranging from 4.7 to 6.2 µM, have also been reported in MMV-Luc cells, a luciferase-expressing reporter cell line used to assess estrogenic activity (Demaegdt et al. 2016; Frizzell et al. 2013). In a study using an integrated *in silico/in vitro* approach, AOH showed structural similarity to estradiol with methylation further enhancing its estrogenic potential (Dellafiora et al. 2018). As part of the 2011 risk assessment of *Alternaria* toxins, the EFSA CONTAM Panel also concluded that AOH and AME may interfere with estrogen receptor-mediated pathways, although mechanisms could not be elucidated due to cell-type specificity and incomplete data (EFSA 2011).

TeA induced luciferase expression at the highest concentration used in the agonistic part of the study, but it was not considered positive based on the OECD TG criteria. Although not identified as an agonist when tested alone, TeA at the highest concentration when co-incubated with E2, elicited an increased luciferase signal response (above that of E2 control alone). ALT, TEN and ATX-I were negative for ER-mediated agonistic effects while none of the tested compounds showed ER-mediated antagonistic effects except for ATX-I which induced a weak but significant inhibition of the relative ER-mediated transcriptional activity. Although the decrease did not reach 50% of the relative transcriptional activity of estradiol alone, the assay was considered positive based on the OECD TG 455 criteria for antagonistic effects. In Ishikawa cells, the perylene quinones ATX-I and alterperylenol have been reported to mediate antiestrogenic activity, measured as expression of alkaline phosphatase (Crudo et al. 2025). Weak ER-antagonistic effects of low potency have been reported in the literature for AOH only when high concentrations were used (Demaegdt et al. 2016).

The E-Screen bioassay was also conducted to assess the effects of *Alternaria* toxins on MCF-7 cell proliferation (Soto et al. 1995). A DNA-content-based assay was employed to estimate cell number/proliferation, which is less susceptible to changes in cellular metabolic activity than commonly used enzymatic metabolism-based assays (Quent et al. 2010). Although a statistically significant increase in MCF-7 cell proliferation was observed for all compounds except TeA, the effect was confined to the highest non-cytotoxic concentration only. In addition, various non-estrogenic substances have been reported to alter MCF-7 cell proliferation, which may also be mediated by signaling pathways other than estrogen receptor activation (Kinnberg 2003). Thus, the results obtained from the E-screen assay should be interpreted as supplementary to the OECD TG 455. In consistence with the results of the OECD compliant ER-transactivation assay, ATX-I induced significant decreases in the E2-induced proliferation of MCF-7 cells at concentrations that were not cytotoxic when tested alone.

For assessing AR-mediated activity, the OECD TG 458 assay has been conducted using the AR EcoScreen cell line. According to the OECD TG, AR-EcoScreen cells may show limited GR crosstalk due to the recognition of shared hormone response elements by AR and GR. Thus, co-incubation with the specific AR antagonist hydroxyflutamide (HF) is recommended to confirm whether the observed agonistic activity is AR-mediated. For compounds that test positive but lack mechanistic information, the response in the presence of HF may also provide indications of potential GR involvement (Rosenmai et al. 2018). In the present study, weak positive effects in line with the OECD criteria have been identified for AOH and AME in two individual experimental runs. Although some variability in the concentrations showing positive effects between independent experiments was observed possibly due to substance degradation in the stock solutions used, all experimental runs were considered weakly positive for AOH and AME. As expected, a marked decrease in luciferase activity was observed for DHT in the presence of HF. Interestingly, significant increases in luciferase activity were still observed for AOH and AME while AOH fulfilled the OECD positivity criteria in the presence of HF, indicating a possible involvement of GR in the minimal inductions that were observed. This was further supported by the true AR-responses calculated according to OECD, after subtracting the response observed in the presence of HF, which were below the 10% threshold for positivity for both AOH and AME. Based on the available evidence, no interaction with the AR was observed for AOH in two independent studies using the androgen-dependent TARM-Luc cell line (Demaegdt et al. 2016; Frizzell et al. 2013). On the other hand, Stypuła-Trębas et al. (2017) suggested a full androgenic response for AOH in a yeast bioassay, although potency was considerably low (Stypuła-Trębas et al. 2017). Frizzell et al. 2013, used mammalian reporter gene assays with cells transfected with steroid hormone receptor constructs, including GR, and no effect was observed for AOH. Our results could be indicative of a possible interaction between AR- and GR-mediated signaling, suggesting that both receptors may play a role in the observed response. Future investigations using systems such as the AR-EcoScreen GR-knockout cell line or the Human Glucocorticoid Receptor Transactivation Assay in accordance with OECD TG 454A (OECD, 2026) may provide additional insights into the potential interaction of AOH and AME with GR-mediated signaling pathways.

No AR antagonistic activity was observed in the EcoScreen assay for AOH, AME, ATX-I, TEN, ALT and TeA at non cytotoxic doses. In a study using stably transfected TARM-Luc cells, AOH has been shown to decrease luciferase activity at high doses. According to the authors, as a similar effect was also observed in progestagen and glucocorticoid responsive cells, the decrease is more likely correlated with cytotoxicity rather than AR mediated antagonism. However, strong decreases in luciferase activity were observed at the dose of 4.84 and 9.68 μM, in the absence of cytotoxicity (Frizzell et al. 2013). Based on Demaegdt et al. (2016) that used the same cell line, AOH exhibited AR mediated antagonistic effects in the absence of cytotoxicity, with an IC_50_ of 3.8 μM. Strong decreases in luciferase activity were also observed in the present study at the dose of 30 μM for AOH and AME and from the dose of 10 μM for ATX-I. However, as cytotoxicity above 20% was also observed at the same doses, the assay is considered overall negative based on the OECD TG 458 criteria (Demaegdt et al. 2016). This suggests that the differences from previous findings may be related to compound-induced cytotoxicity, under the conditions used in the present study.

No effects on estradiol or testosterone production were observed for *Alternaria* toxins when assessed in the OECD TG 456 assay in H295R cells. Although ATX-I induced a statistically significant decrease in testosterone production at a non-cytotoxic concentration, the effect did not exceed the 1.5-fold threshold and the assay was considered overall negative based on the OECD criteria. In another study using the same cell line, AOH has been shown to increase estradiol synthesis at the highest concentration used (3.87 μM) and to induce mRNA expression of steroidogenesis-related genes (Frizzell et al. 2013). The authors further examined the proteomic response of H295R cells to AOH, and identified regulated proteins involved in the early-stage steroid biosynthesis and in C21-steroid hormone metabolism (Kalayou et al. 2014). On the other hand, AOH and AME administration to pig granulosa cells, decreased the level of progesterone secretion without altering the expression of steroidogenesis-related genes, while TeA showed no effect on progesterone secretion in the same cell system (Tiemann et al. 2009). In a more recent study, AOH enhanced DHEA-driven local steroidogenesis in human normal prostate epithelial cells and prostate cancer cells, affecting steroid hormone production and modulating the expression of key steroidogenic markers (Urbanek et al. 2023).

The present study aimed to address existing gaps in the EAS mediated activity of *Alternaria* toxins using validated assays conducted according to OECD TGs and well characterized compounds. Testing requirements for assessing the EAS modalities are fulfilled by either the OECD Test Guideline 443 extended one-generation reproductive toxicity study (OECD 2025a), or the OECD TG 416 two-generation reproductive toxicity study (OECD 2001). In the case of an incomplete *in vivo* dataset, a tiered testing strategy is applied, starting with the OECD *in vitro* assays assessing EAS- mediated activity. These include the OECD TG 455, OECD TG 458 and OECD TG 456 assays. If negative, *in vitro* testing is followed by the Hershberger OECD TG 441 assay (OECD 2009) for the assessment of A modality and the Uterotrophic OECD TG 440 assay (OECD 2007) for the assessment of E modality. In the case of negative results, it may be concluded that ED criteria are not met for E modality. The ToxCast ER model value (Browne et al. 2015) may be also used to conclude that the ED criteria are not met for E modality. When concern is identified based on the assessment of endocrine activity, further testing may be required. The current results on the *in vitro* EAS-mediated activity of *Alternaria* toxins, provide valuable data that can be interpreted within the current regulatory framework to support ED assessments, identify potential concerns and prioritize *Alternaria* compounds for further testing.

## Statements and Declarations

The authors declare that they have no conflict of interest.

## Acknowledgments

The European Partnership for the Assessment of Risks from Chemicals (PARC) has received funding from the European Union’s Horizon Europe research and innovation program under Grant Agreement No. 101057014 and has received co-funding of the authors’ institutions. Views and opinions expressed are, however, those of the author(s) only and do not necessarily reflect those of the European Union or the Health and Digital Executive Agency. Neither the European Union nor the granting authority can be held responsible for them.

The authors gratefully acknowledge Dr. Mitsuru Iida, developer of the AR-EcoScreen cell line, for kindly providing the cell line used in this study.

## Supplementary Information

**SI-1:** Results of the agonistic effects of AOH and AME in the AR STTA from two individual experimental runs

| AOH Agonism |  |  |  |  |
| --- | --- | --- | --- | --- |
| Compound/Experimental run/concentration | % viability | SD | % RTA | SD |
| AOH, Exp1, 0.1µM | 111.31 | 3.20 | 1.11 | 3.31 |
| AOH, Exp1, 0.3µM | 123.28 | 10.20 | 3.82 | 1.70 |
| AOH, Exp1, 1µM | 116.90 | 10.97 | 2.69 | 1.30 |
| AOH, Exp1, 3µM | 117.36 | 6.54 | **10.86** | 0.97 |
| AOH, Exp1, 10µM | 115.88 | 10.56 | **19.38** | 3.32 |
| AOH, Exp1, 30µM | 48.22 | 2.68 | 1.82 | 0.47 |
| AOH, Exp1, 60µM | 6.60 | 1.60 | -19.19 | 0.35 |
| AOH, Exp2, 0.1µM | 113.52 | 13.70 | -0.26 | 1.87 |
| AOH, Exp2, 0.3µM | 119.87 | 4.46 | -2.08 | 1.17 |
| AOH, Exp2, 1µM | 112.57 | 6.26 | -0.80 | 0.29 |
| AOH, Exp2, 3µM | 114.38 | 12.07 | 1.19 | 1.19 |
| AOH, Exp2, 10µM | 105.09 | 12.43 | 7.93 | 1.29 |
| AOH, Exp2, 30µM | 93.24 | 5.01 | **15.40** | 1.61 |
| AOH, Exp2, 60µM | 23.47 | 2.75 | -9.22 | 0.49 |
| AOH, Exp3, 0.1µM | 86.08 | 11.6 | 3.11 | 0.42 |
| AOH, Exp3, 0.3µM | 88.32 | 13.37 | 2.67 | 0.40 |
| AOH, Exp3, 1µM | 91.87 | 10.50 | 3.72 | 0.85 |
| AOH, Exp3, 3µM | 89.02 | 9.55 | 4.70 | 0.53 |
| AOH, Exp3, 10µM | 87.66 | 12.40 | **10.30** | 0.08 |
| AOH, Exp3, 30µM | 44.79 | 6.49 | 11.90 | 1.57 |

**In bold:** Positive response in line with OECD TG criteria
| AME Agonism |  |  |  |  |
| --- | --- | --- | --- | --- |
| Compound/Experimental run/concentration | % viability | % SD | % RTA | % SD |
| AME, Exp1, 0.1µM | 124.24 | 17.54 | 4.67 | 0.21 |
| AME, Exp1, 0.3µM | 129.26 | 11.22 | 3.68 | 1.02 |
| AME, Exp1, 1µM | 126.27 | 5.97 | 4.31 | 1.03 |
| AME, Exp1, 3µM | 132.74 | 4.32 | 6.01 | 0.96 |
| AME, Exp1, 10µM | 129.07 | 16.01 | **20.62** | 1.67 |
| AME, Exp1, 30µM | 85.21 | 9.93 | **20.64** | 6.63 |
| AME, Exp1, 60µM | 41.05 | 3.76 | 16.32 | 2.14 |
| AME, Exp2, 0.1µM | 92.56 | 10.21 | -1.04 | 0.76 |
| AME, Exp2, 0.3µM | 95.63 | 6.16 | -1.14 | 0.61 |
| AME, Exp2, 1µM | 100.59 | 0.77 | -0.18 | 1.24 |
| AME, Exp2, 3µM | 98.78 | 11.22 | -0.07 | 0.89 |
| AME, Exp2, 10µM | 100.64 | 2.89 | 2.79 | 1.00 |
| AME, Exp2, 30µM | 97.62 | 3.66 | **10.86** | 1.65 |
| AME, Exp2, 60µM | 54.74 | 11.80 | 7.67 | 2.26 |
| AME, Exp3, 0.1µM | 101.61 | 15.69 | 1.21 | 0.52 |
| AME, Exp3, 0.3µM | 107.55 | 16.30 | 2.22 | 0.62 |
| AME, Exp3, 1µM | 106.03 | 12.76 | 2.08 | 0.51 |
| AME, Exp3, 3µM | 108.32 | 11.10 | 3.41 | 1.80 |
| AME, Exp3, 10µM | 98.30 | 12.04 | **10.00** | 2.73 |
| AME, Exp3, 30µM | 86.00 | 13.46 | **18.91** | 1.90 |

**SI-2:**
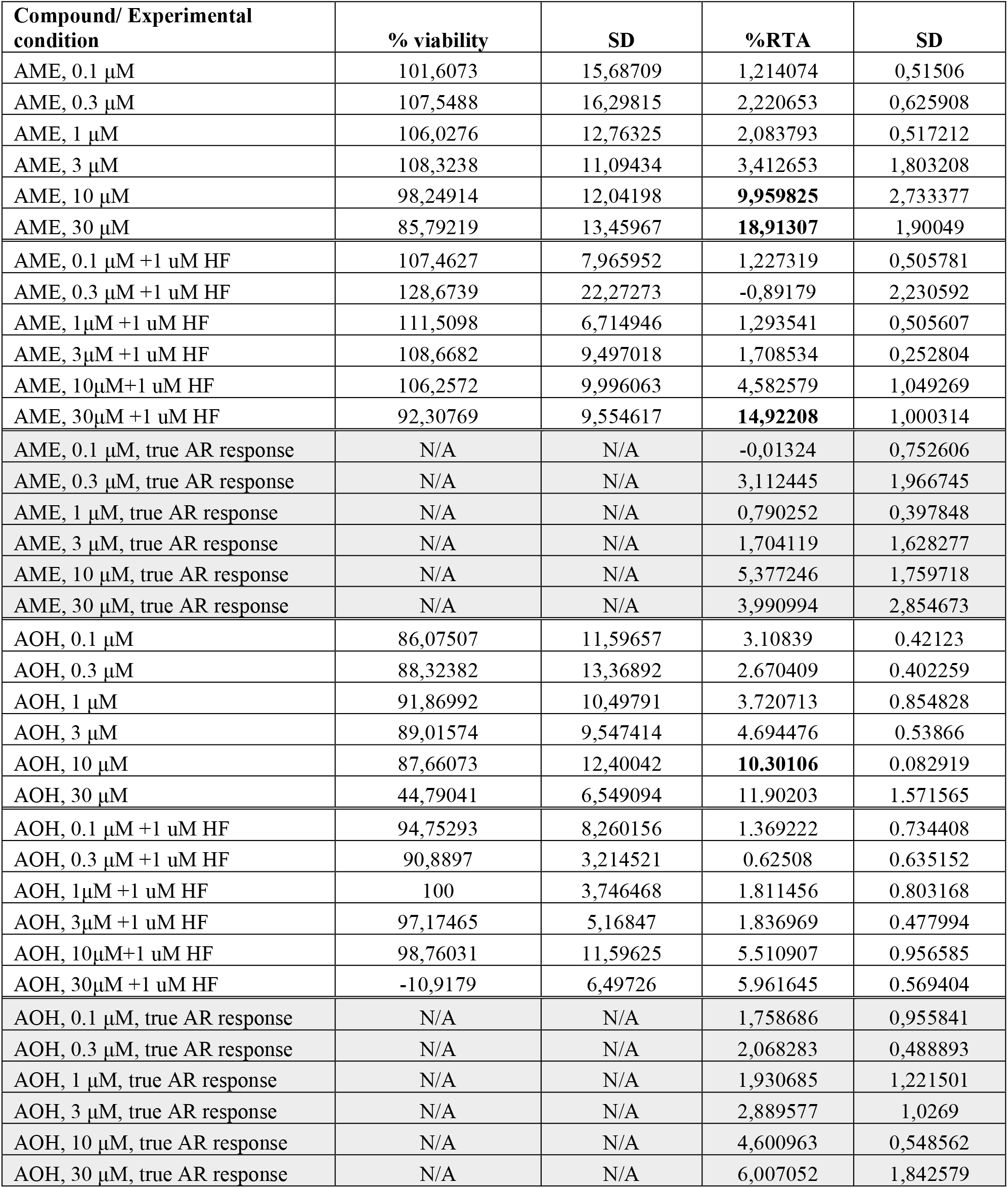
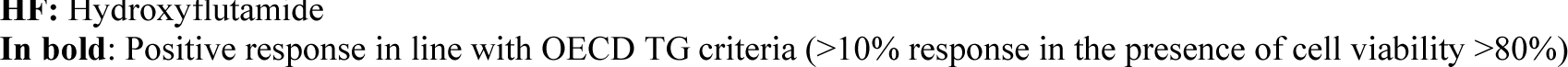
True response values for AME and AOH after co-incubation with the specific AR antagonist hydroxyflutamide (HF)

**In bold:** Positive response in line with OECD TG criteria **SI-2:** True response values for AME and AOH after co-incubation with the specific AR antagonist hydroxyflutamide (HF)
| Compound/ Experimental condition | % viability | SD | %RTA | SD |
| --- | --- | --- | --- | --- |
| AME, 0.1 μM | 101,6073 | 15,68709 | 1,214074 | 0,51506 |
| AME, 0.3 μM | 107,5488 | 16,29815 | 2,220653 | 0,625908 |
| AME, 1 μM | 106,0276 | 12,76325 | 2,083793 | 0,517212 |
| AME, 3 μM | 108,3238 | 11,09434 | 3,412653 | 1,803208 |
| AME, 10 μM | 98,24914 | 12,04198 | **9,959825** | 2,733377 |
| AME, 30 μM | 85,79219 | 13,45967 | **18,91307** | 1,90049 |
| AME, 0.1 μM +1 uM HF | 107,4627 | 7,965952 | 1,227319 | 0,505781 |
| AME, 0.3 μM +1 uM HF | 128,6739 | 22,27273 | -0,89179 | 2,230592 |
| AME, 1μM +1 uM HF | 111,5098 | 6,714946 | 1,293541 | 0,505607 |
| AME, 3μM +1 uM HF | 108,6682 | 9,497018 | 1,708534 | 0,252804 |
| AME, 10μM+1 uM HF | 106,2572 | 9,996063 | 4,582579 | 1,049269 |
| AME, 30μM +1 uM HF | 92,30769 | 9,554617 | **14,92208** | 1,000314 |
| AME, 0.1 μM, true AR response | N/A | N/A | -0,01324 | 0,752606 |
| AME, 0.3 μM, true AR response | N/A | N/A | 3,112445 | 1,966745 |
| AME, 1 μM, true AR response | N/A | N/A | 0,790252 | 0,397848 |
| AME, 3 μM, true AR response | N/A | N/A | 1,704119 | 1,628277 |
| AME, 10 μM, true AR response | N/A | N/A | 5,377246 | 1,759718 |
| AME, 30 μM, true AR response | N/A | N/A | 3,990994 | 2,854673 |
| AOH, 0.1 μM | 86,07507 | 11,59657 | 3.10839 | 0.42123 |
| AOH, 0.3 μM | 88,32382 | 13,36892 | 2.670409 | 0.402259 |
| AOH, 1 μM | 91,86992 | 10,49791 | 3.720713 | 0.854828 |
| AOH, 3 μM | 89,01574 | 9,547414 | 4.694476 | 0.53866 |
| AOH, 10 μM | 87,66073 | 12,40042 | **10.30106** | 0.082919 |
| AOH, 30 μM | 44,79041 | 6,549094 | 11.90203 | 1.571565 |
| AOH, 0.1 μM +1 uM HF | 94,75293 | 8,260156 | 1.369222 | 0.734408 |
| AOH, 0.3 μM +1 uM HF | 90,8897 | 3,214521 | 0.62508 | 0.635152 |
| AOH, 1μM +1 uM HF | 100 | 3,746468 | 1.811456 | 0.803168 |
| AOH, 3μM +1 uM HF | 97,17465 | 5,16847 | 1.836969 | 0.477994 |
| AOH, 10μM+1 uM HF | 98,76031 | 11,59625 | 5.510907 | 0.956585 |
| AOH, 30μM +1 uM HF | -10,9179 | 6,49726 | 5.961645 | 0.569404 |
| AOH, 0.1 μM, true AR response | N/A | N/A | 1,758686 | 0,955841 |
| AOH, 0.3 μM, true AR response | N/A | N/A | 2,068283 | 0,488893 |
| AOH, 1 μM, true AR response | N/A | N/A | 1,930685 | 1,221501 |
| AOH, 3 μM, true AR response | N/A | N/A | 2,889577 | 1,0269 |
| AOH, 10 μM, true AR response | N/A | N/A | 4,600963 | 0,548562 |
| AOH, 30 μM, true AR response | N/A | N/A | 6,007052 | 1,842579 |

**HF:** Hydroxyflutamide
**In bold:** Positive response in line with OECD TG criteria (>10% response in the presence of cell viability >80%)

**SI-3:**
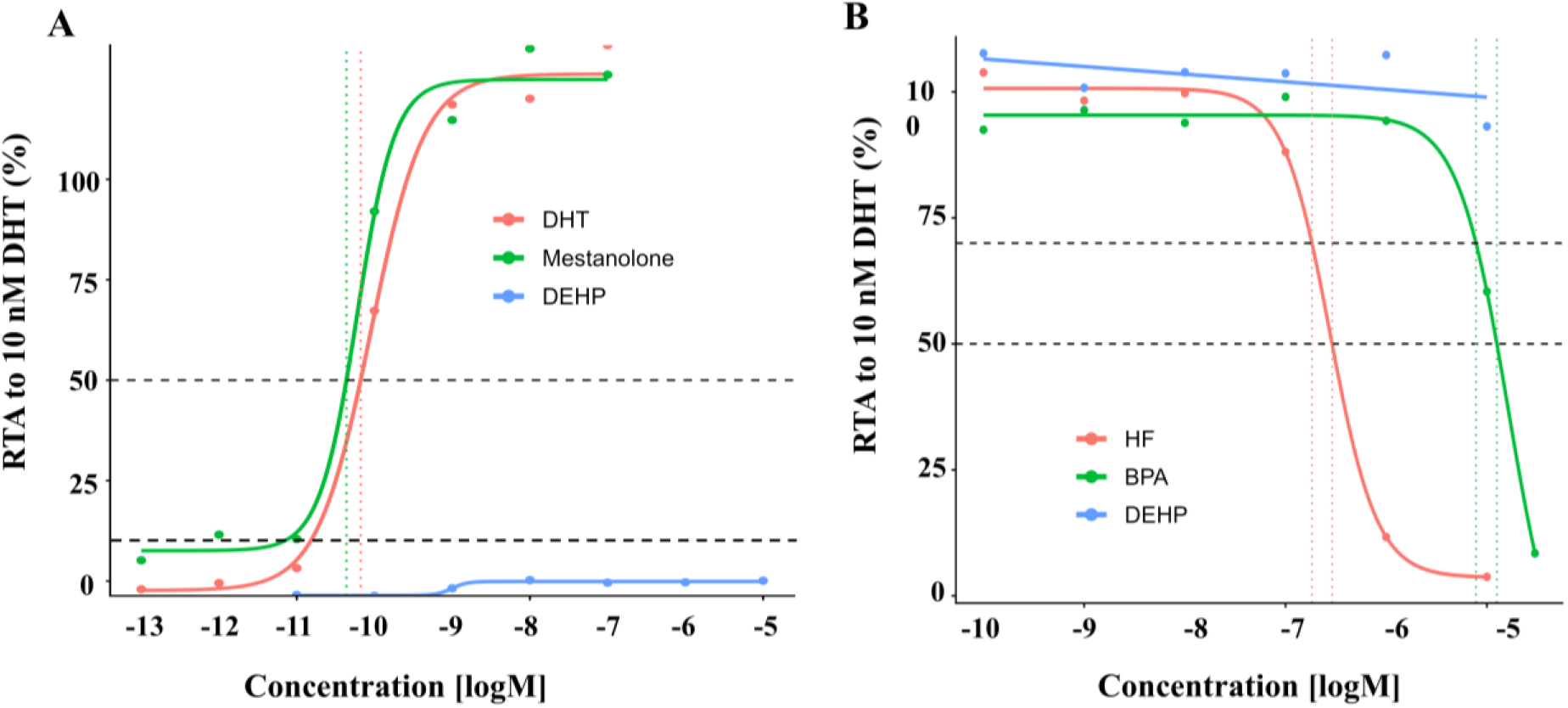
Concentration response curves for reference/proficiency compounds for the agonistic (A) and antagonistic (B) part of the AR STTA (AR-EcoScreen, OECD TG 458)

**SI-4:** Decision matrix of the steroidogenesis assay (OECD TG 456)

Estradiol
|  | Run 1 | Run 2 | Run 3 | Final call |
| --- | --- | --- | --- | --- |
| AOH | negative | negative | negative | negative |
| AME | negative | equivocal | negative | negative |
| ALT | negative | negative | negative | negative |
| ATX-I | equivocal | negative | negative | negative |
| TeA | negative | negative | negative | negative |
| TEN | negative | negative | negative | negative |

Testosterone
|  | Run 1 | Run 2 | Run 3 | Final call |
| --- | --- | --- | --- | --- |
| AOH | equivocal | negative | negative | negative |
| AME | negative | equivocal | negative | negative |
| ALT | negative | negative | negative | negative |
| ATX-I | equivocal | negative | negative | negative |
| TeA | negative | negative | negative | negative |
| TEN | negative | negative | negative | negative |

**SI-5.**
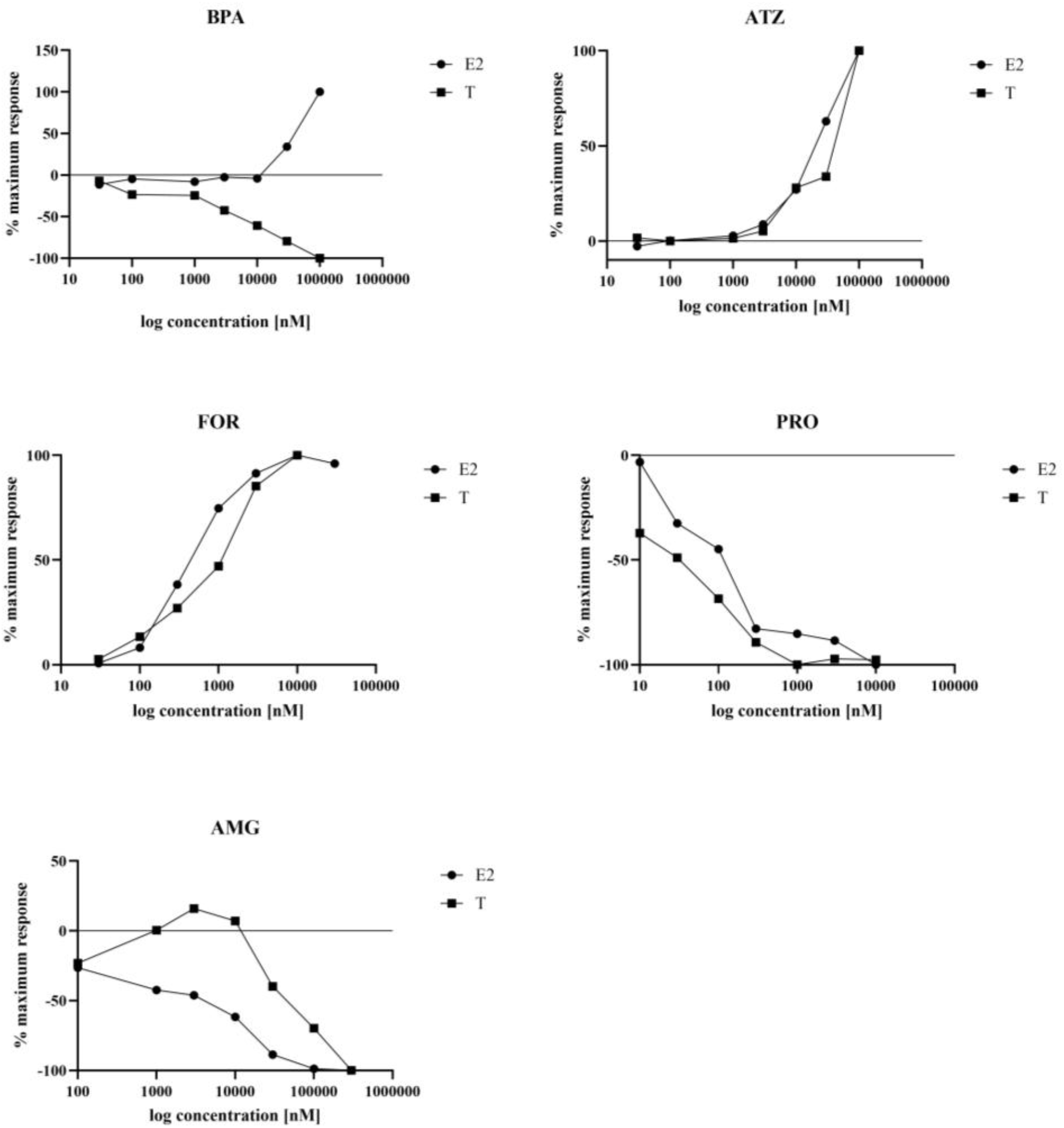
Maximum response curves from the steroidogenesis proficiency test

